# Mechanisms underlying reshuffling of visual responses by optogenetic stimulation in mice and monkeys

**DOI:** 10.1101/2022.07.13.499597

**Authors:** A. Sanzeni, A. Palmigiano, T.H. Nguyen, J. Luo, J.J. Nassi, J.H. Reynolds, M.H. Histed, K.D. Miller, N. Brunel

## Abstract

The ability to optogenetically perturb neural circuits opens an unprecedented window into mechanisms governing circuit function. We analyzed and theoretically modeled neuronal responses to visual and optogenetic inputs in mouse and monkey V1. In both species, optogenetic stimulation of excitatory neurons strongly modulated the activity of single neurons, yet had weak or no effects on the distribution of firing rates across the population. Thus, the optogenetic inputs reshuffled firing rates across the network. Key statistics of mouse and monkey responses lay on a continuum, with mice/monkeys occupying the low/high rate regions, respectively. We show that neuronal reshuffling emerges generically in randomly connected excitatory/inhibitory networks, provided the coupling strength (combination of recurrent coupling and external input) is sufficient that powerful inhibitory feedback cancels the mean optogenetic input. A more realistic model, distinguishing tuned visual vs. untuned optogenetic input in a structured network, reduces the coupling strength needed to explain reshuffling.

## Introduction

Since their introduction nearly two decades ago, optogenetic methods have revolutionized neuroscience (Kim et al., 2017). In particular, optical stimulation of specific cell types, in conjunction with electrophysiological or optical recordings of neuronal activity, can be used to probe network operating regime and test predictions from theoretical models. Successful examples include testing the ‘paradoxical effect’ in mouse cortex in inhibition-stabilized networks (Ozeki et al., 2009; Sanzeni et al., 2020; Tsodyks and Markram, 1997), and testing predictions of attractor models in both mouse cortex (Inagaki et al., 2019) and the fly central complex (Kim et al., 2017). Optogenetics, in conjunction with theoretical modeling, can also provide information about the structure of synaptic connectivity (*e.g*. Palmigiano et al., 2020). For instance, activation of single neurons in V1 suppresses other neurons, with a magnitude depending on the distance between their preferred orientations (Chettih and Harvey, 2019), consistent with a network with strong orientation specificity of excitatory-inhibitory (E-I) connectivity (Ko et al., 2011; Sadeh and Clopath, 2020).

To probe the operating regime of sensory cortex (Ahmadian and Miller, 2021) and the mechanisms that shape its responses to its inputs, we analyze electrophysiological recordings from both mouse (Histed, 2018) and monkey (Nassi et al., 2015) primary visual cortex (V1) during presentation of visual stimuli of various contrasts and/or optogenetic stimulation of E neurons. The optogenetic stimulation leads to a broad distribution of firing rate changes in single neurons, with some neurons being strongly excited, while others are strongly suppressed. The typical firing rate changes in single neurons are much larger than the changes in mean response of the population, which are only significant for lower visual contrasts in mice. In fact, the entire distribution of firing rates is largely unaffected by optogenetic stimulation (significant changes seen only at low visual contrast in mice), even though the variance of firing rate changes is comparable to the variance of rates prior to optogenetic stimulation. Thus, optogenetic stimulation of E neurons leads to a reshuffling of firing rates across the network, leaving invariant the whole distribution. We find a continuum from mice at low contrast to mice at high contrast to monkeys at low contrast to monkeys at high contrast, along which firing rates increase and the optogenetically-induced changes in the mean and distribution of firing rates shrink to zero.

We explore the mechanisms of such a reshuffling using theoretical analysis and modeling. We first examine randomly connected networks of E and I neurons. We find that such networks reproduce all the above mentioned features, provided the strengths of recurrent coupling and external input are strong enough, the network is inhibition dominated, and opsin-induced currents in E neurons are sufficiently heterogeneous and weaker in the mean than visually-induced currents. The continuity from mice to monkeys can be understood if mice are in a more weakly-coupled regime than monkeys, which can arise from weaker recurrent weights and/or weaker external inputs. We also investigate structured network models with connectivity dependent on spatial distance and orientation tuning, which can incorporate the difference between tuned visual and untuned optogenetic input. We find that reshuffling can arise in structured networks with weaker effective coupling and stronger optogenetic input than in the unstructured randomly connected model. These results provide new insights into the operating regime of sensory cortex and how this may differ across different mammalian species.

## Results

### Heterogeneous optogenetic modulation of visual responses in mice and monkeys

We analyzed recordings obtained in V1 of mice (data from Histed, 2018) and monkeys (data from Nassi et al., 2015). These experiments shared a number of common features (see *Methods* for details): visual stimuli were gratings of different contrasts ([0 – 90]% in mice and [0 – 99]% in monkeys); virally expressed opsins were used to optogenetically stimulate pyramidal cells. Both datasets consist of neural spiking responses recorded extracellularly in awake animals to combinations of visual and optogenetic stimuli. There are also a few differences between the recordings in (Histed, 2018) and (Nassi et al., 2015). In monkeys, visual stimuli were drifting sinusoidal gratings centered within the receptive field and matched to each recorded unit’s preferred size, orientation, and spatial and temporal frequencies. In mice, on the other hand, multiple neurons were recorded simultaneously and there was no systematic relationship between the visual stimulus and preferences of the recorded neurons.

Both in mice and monkeys, as previously observed (Histed, 2018; Nassi et al., 2015), single neuron responses to visual stimuli were strongly modulated by optogenetic stimuli, in a highly heterogeneous fashion (Fig. 1A-B). Specifically, responses to visual stimuli were typically either enhanced by the laser at all contrasts, suppressed by the laser at all contrasts, or enhanced at low contrast and suppressed at high contrast (Fig. 1A-B); the fourth possibility, suppressed at low contrast and enhanced at high contrast, was not observed. These single neuron responses might underlie different scenarios at the population level (Fig. 1C). (i) Optogenetic stimulation could significantly increase the mean activity in the network, while broadening the distribution of single neuron rates. This scenario is what one would expect naively in experiments in which optogenetic stimulation targets E neurons. (ii) Optogenetic stimulation could broaden the distribution of rates, but a large number of suppressed cells could lead to a weak mean increase of the population activity. (iii) Optogenetic stimulation could affect only weakly the distribution of rates. Which of these scenarios better represents the effect of optogenetic stimulation on population activity? Do these effects change with contrast and species?

**Figure 1:**
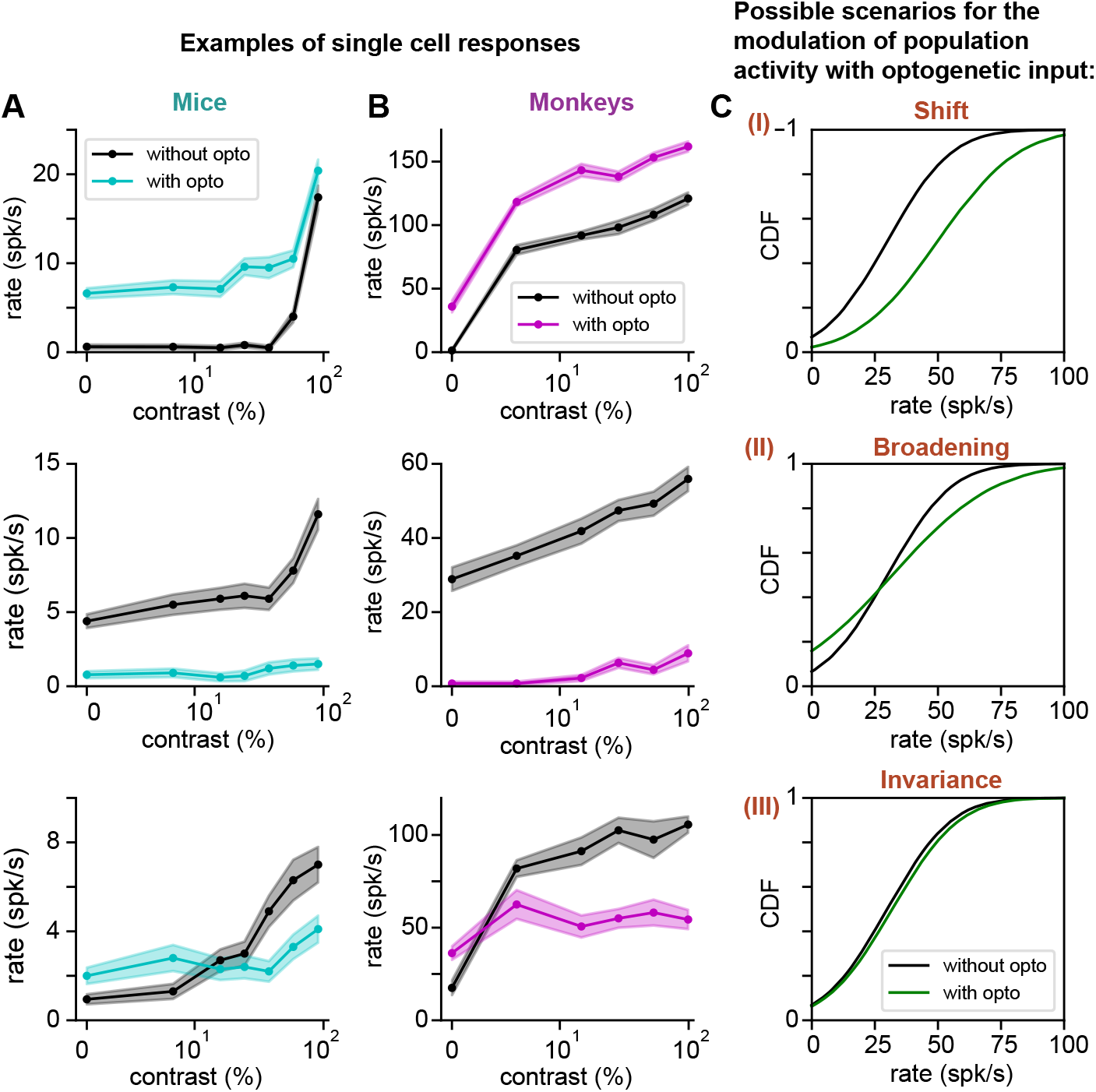
Optogenetic modulation of visual responses in mice and monkeys: single neuron examples and potential scenarios at the population level. (**A**) Examples of single neuron responses to visual stimuli in mice, without (black) and with (colored) optogenetic stimulation of E neurons (here and in the following figure, lines represent means, shaded regions are mean±sem). (**B**) As in **A** but in monkeys. In both species, some neurons strongly increase their rates at all contrasts (top row); others are strongly suppressed (center row); and others exhibit rate increase at low contrast, but suppression at high contrast (bottom row). Note that the same color code is used in all the following figures. (**C**) Possible scenarios of how optogenetic stimulation of E neurons might modify the distribution of firing rates in the network (see text).

### Optogenetic stimulation reshuffles visual responses in mice and monkeys

To answer the above mentioned questions, we analyzed the statistics of population responses to visual and optogenetic stimuli (Fig. 2). In both mice and monkeys, mean population activity increased monotonically with the contrast of visual stimuli, but firing rates in mice were significantly lower than in monkeys. The mean visual response in mice ranged from 6 ± 0.9 spk/s (mean±sem) at 0% contrast to 16.7 ± 2.0 spk/s at high contrast, while in monkeys it ranged from 20.2 ± 2.0 spk/s to 72.7 ± 6.7 spk/s (Fig. 2A). In both species, visual stimuli of fixed contrast generated a broad distribution of rates (Fig. 2B), and increasing contrast significantly shifted distributions toward higher rates (Fig. 2C–D). Surprisingly, optogenetic stimulation had much weaker effects at the population level (Fig. 2A–D). In mice, where all cells were recorded regardless of their feature preference, the mean and standard deviation of the population activity were significantly modulated by the optogenetic stimulation only for contrasts lower than 55% (Welch’s t–test, p–value<0.05) and 90%(f–test, p–value<0.05), respectively. In monkeys, where cells were only recorded with stimuli matching their feature preferences, the mean population activity was not significantly modulated at any contrast (Welch’s t–test, p–value>0.05) while the modulation of the standard deviation was significant only at contrasts lower than 12% (f-test, p–value<0.05). Even more strikingly, optogenetic stimuli weakly affected the whole distribution of rates in the network (Fig. 2D). In mice, the distribution of rates was significantly affected only at low contrast (< 20%, KS test, p–value<0.05), for which optogenetic stimuli induced a weak but statistically significant shift of the distribution toward higher rates. In monkeys, the distribution of rates was not significantly affected by optogenetic stimulation at any contrast tested (KS test, p–value>0.05).

**Figure 2:**
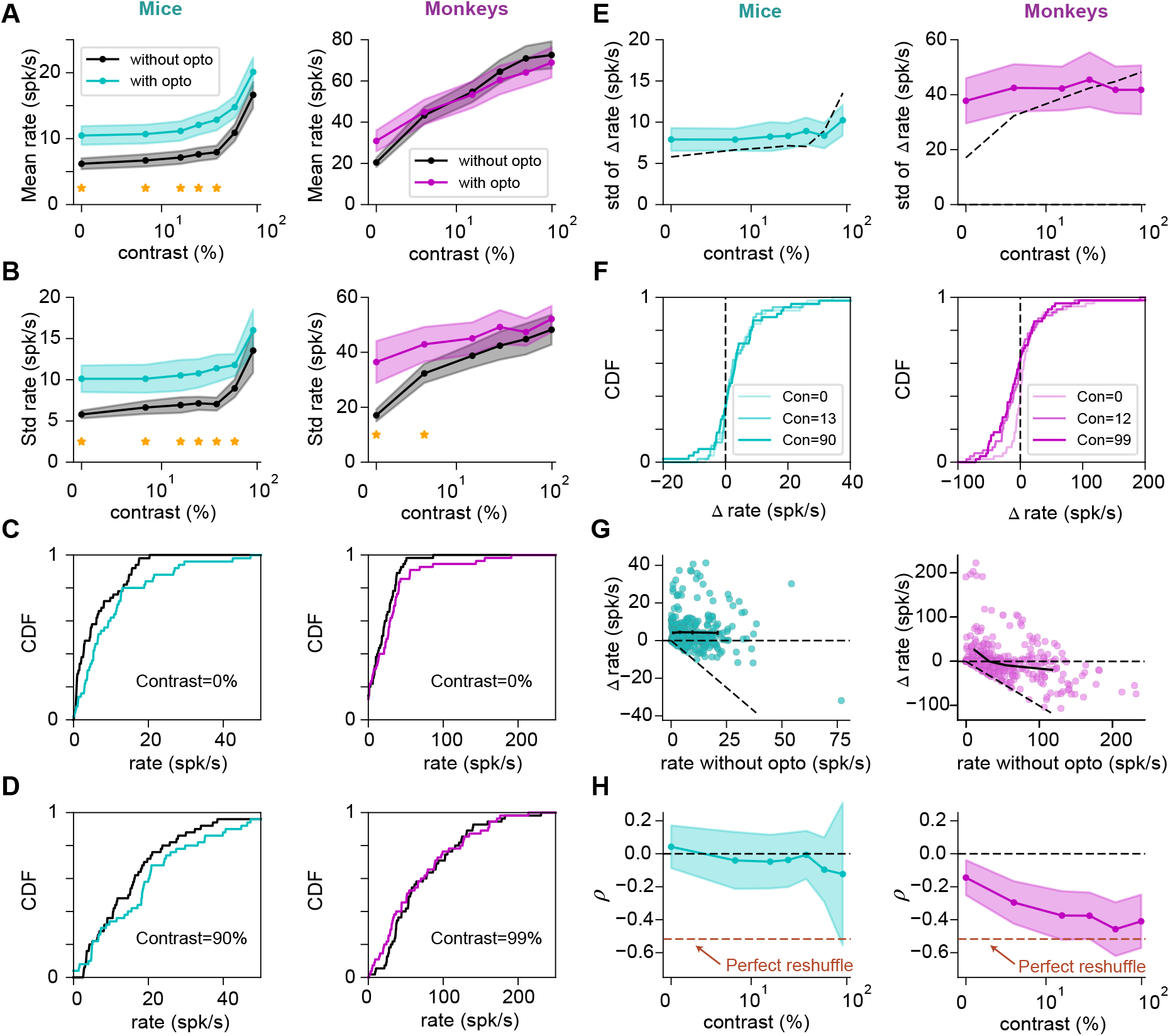
Effects of optogenetic stimulation on population statistics in mice and monkeys. (**A**) Population mean response to visual and optogenetic stimuli. Optogenetic modulation caused no significant change in mean rates except in mice at low contrast (< 55%, Welch’s t–test, p–value<0.05). (**B**) Standard deviation of the distribution of firing rates across the population. In both species, the standard deviation of firing rates is increased significantly by light only at low contrast (in mice, < 90%; In monkeys, < 12%; f-test, p–value<0.05). (**C-D**) Cumulative distribution function (CDF) of firing rates at low (**C**) and high (**D**) contrasts of visual stimuli, with (colored) and without (black) optogenetic stimuli. Optogenetic modulation was only significant for mice at low contrast (6% and 12%, KS test, p–value<0.05). (**E**) Standard deviation of the change in firing induced by optogenetic stimulation as a function of contrast of the visual stimulus (colored lines). In both species, the standard deviation of the change in firing rate was larger than the mean and of the same order of the standard deviation of rates without the laser (dashed black lines, same as black lines in B). (**F**) Cumulative distribution function of changes in firing rate (Δ*r*) induced by optogenetic stimulation at three example contrasts (indicated by transparency levels). In both species, and at all contrasts, optogenetic stimuli generated a broad distribution of firing rate changes, with a large fraction of suppressed cells. (**G**) Change in firing rate induced by optogenetic stimulation as a function of baseline activity (i.e. activity without optogenetic stimuli). Each cell is represented once for every value of contrast of the visual stimuli. Continuous black lines represent mean Δ*r* in quartiles of the distribution of rates without optogenetic stimuli. (**H**) Normalized covariance of baseline rates and change in rates induced by optogenetic stimuli (*ρ* =Cov(*r*, Δ*r*)/Var(Δ*r*)). This ratio decreases with contrast in both species and, in monkeys, it approaches the value −0.5 which, as described in the text, is required to obtain perfect invariance of the distribution of rates.

The weak optogenetic modulation observed at the population level was not due to small changes in single neuron firing. In fact, both in mice and monkeys, distributions of optogenetic modulations of single neuron firing rates were broad, with standard deviation comparable to or larger than the standard deviation of the rates without the optogenetic stimulus (Fig. 2E–F). Furthermore, unlike what was found in response to visual stimuli (Fig. 2C–D), the mean changes in rate due to optogenetic stimulation were characterized by a mean that was considerably smaller than the standard deviation of the rate changes (Fig. 2A, E). Specifically, combining all contrasts, changes in rates due to optogenetic input in mice and monkeys were 4.2 ± 8.9 spk/s (mean±std) and −0.7± 43 spk/s (mean±std), respectively. The distribution of optogenetic responses was contrast invariant in mice (KS test, p-value>0.05). In monkeys, on the other hand, optogenetic responses observed at zero contrast followed a distribution that was significantly different (KS test, p-value<0.05) from those at higher contrasts, which were not significantly different from one another. Furthermore, a significant fraction of cells were suppressed (e.g. at high contrast, 16/55 in mice and 37/50 in monkeys, respectively).

In sum, the response statistics characterized here suggest that optogenetic stimuli *reshuffled* visual responses in mice and monkeys: they strongly modulated single neuron responses, but overall led to a weak modulation of population activity.

How can optogenetic stimulation leave the distribution of rates unchanged, while at the same time leading to large changes in firing rates of individual neurons (Fig. 2B)? If optogenetic responses were independent of baseline firing, as one might expect intuitively, optogenetic stimuli would significantly broaden the distribution of visual responses. The rate variance with 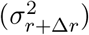 or without 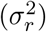 optogenetic stimuli are related by 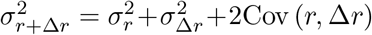 (where “Cov” is covariance). Thus, independence of *r* and Δ*r* would imply the rate variance is increased by 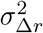, which as we have seen (Fig. 2E) is comparable in size to 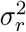. If the distribution of rates is not modified by optogenetic stimuli, then 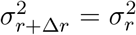, which requires a negative covariance with value 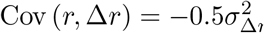.

The relationship between change in rate induced by laser Δ*r* and rate without laser *r* is illustrated in Fig. 2G for both mice and monkeys. In mice, the two are weakly anticorrelated (combining all contrasts, the Pearson correlation coefficient was −0.06), consistent with a significant increase in the width of the rate distribution with optogenetic stimulation, while in monkeys, there is a strong anticorrelation between Δ*r* and *r* (combining all contrasts, the Pearson correlation coefficient was −0.38) – neurons with higher (lower) baseline rates were more likely to be suppressed (excited) by optogenetic stimuli. To quantify the degree to which this anticorrelation leaves the distribution of firing rates invariant, we define a normalized covariance

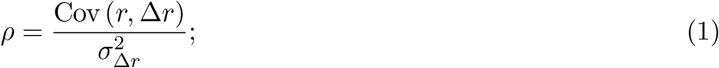

so that *ρ* is equal to −0.5 when optogenetic stimuli leave invariant the distribution of rates. *ρ* is plotted as a function of visual contrast in Fig. 2H. In mice, the normalized covariance is never significantly different from zero, though there is a trend toward a negative covariance at the highest contrasts. In monkeys, it is significantly negative at all contrasts and close to −0.5 at high contrast, consistent with invariance of the distribution of rates and consequent reshuffling of rates by optogenetic stimulation.

The analysis shown so far shows common trends in mice and monkeys but also significant quantitative differences. We wondered whether these quantitative differences might at least partly be explained by the differences between the firing rates observed in the two animals. Therefore, we plotted dimensionless variables characterizing the effect of optogenetic stimuli on the statistics of network activity in the presence of a given visual stimulus as a function of the average firing rate induced by that visual stimulus. Remarkably, Figs. 3A–B show that curves traced by monkey data seem to be the continuation of those for mouse data, with mice occupying the low rate region and monkeys the high rate region of underlying common curves. In particular, in both species, the mean of Δ*r* relative to its standard deviation decreases as a function of contrast until it becomes negligible at high firing rate. The normalized covariance *ρ* also decreases with firing rate, becoming closer to −0.5 at high rate.

**Figure 3:**
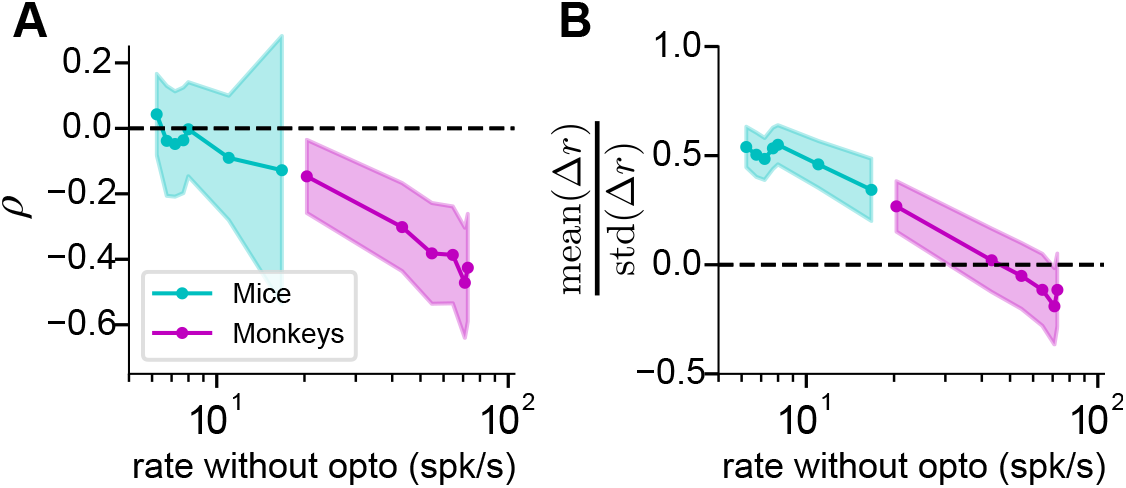
Optogenetic responses in mice and monkeys as a function of average visual response. (**A**) Normalized covariance (Eq. (1)) as a function of average visual response (different points correspond to different contrasts). (**B**) Ratio of mean over standard deviation of the change in rate, as a function of average visual response. Data in both animals seem to lie on the same underlying curves, with data in mice/monkeys covering the low/high rate regions, respectively.

### Network reshuffling emerges in strongly coupled network models

To understand the mechanisms underlying reshuffling of visual responses by optogenetic stimuli, as well as the observed differences between mice and monkeys, we investigated the response properties of a mathematical model of cortical circuits. We analyzed a network of randomly connected E and I rate neurons (Harish and Hansel, 2015; Kadmon and Sompolinsky, 2015; Sompolinsky et al., 1988). In this unstructured model, the single neuron response function (firing rate vs input current) was taken to be the f-I curve of leaky integrate-and-fire neurons driven by white noise (Amit and Brunel, 1997; Amit and Tsodyks, 1991; Siegert, 1951), which is supralinear at lower mean input and becomes sublinear at high input due to saturation, but the results presented here do not depend on this specific choice (see supplementary information, Fig. S1). Visual stimuli are described by external inputs to E and I units, whose strength is drawn from a Gaussian distribution (see details in *Methods*). Optogenetic inputs are described by an additional input to E cells, whose strength is drawn from a log-normal distribution.

Through numerical simulations and analytical calculations, we found that this network model reproduces the main features of the experimental data provided that recurrent connectivity and visual input are strong enough, and the distribution of opsin expression is heterogeneous enough. In particular, we first focus on the dependence of optogenetic responses on the network’s effective synaptic coupling strength (referred to below as coupling strength), which is defined as the synaptic strength times the sensitivity of the postsynaptic cell to synaptic input (Ahmadian and Miller, 2021). This sensitivity is the cell’s gain, i.e. the slope of its input/output function. In the supralinear portion of the input/output function, this increases with network activation and thus with the strength of visual input.

Example network simulations for weak and strong coupling are shown in Fig. 4B-G. In the absence of optogenetic stimulation, and consistent with previous theoretical studies (Amit and Brunel, 1997; Roxin et al., 2011; Sanzeni et al., 2022; van Vreeswijk and Sompolinsky, 1996), the distribution of rates is narrow for weak coupling and broad for strong coupling. Responses to optogenetic stimulation strongly depend on the strength of coupling. For weak coupling, optogenetic stimulation generates a significant shift of the distribution of rates (Fig. 4B). This leads to a broad distribution of optogenetic responses, characterized by a positive mean and a lack of significantly suppressed cells (Fig. 4C). Moreover, single neuron responses to optogenetic stimulation are positively correlated with baseline activity (Fig. 4D, Pearson correlation coefficients are 0.1 and 0.6 for E and I cells, respectively). This positive correlation follows from the supralinear shape of the transfer function at low rates. This effect is more pronounced in I cells because, with respect to E cells, they have a shorter membrane time constant and hence a larger supralinear region (Sanzeni et al., 2020). The response structure observed in weakly coupled networks matches naive expectations but strongly differs from experimental observations. When coupling is strong, on the other hand, optogenetic stimulation does not significantly modify the distribution of rates (Fig. 4E). It produces a broad distribution of rate changes, with a mean that is close to zero and a large fraction of suppressed cells (Fig. 4F). In this regime, single neuron responses to optogenetic stimulation are negatively correlated with baseline activity (Fig. 4G, Pearson correlation coefficient are −0.3 and −0.2 for E and I cells, respectively). These response properties are strikingly similar to experimental observations.

**Figure 4:**
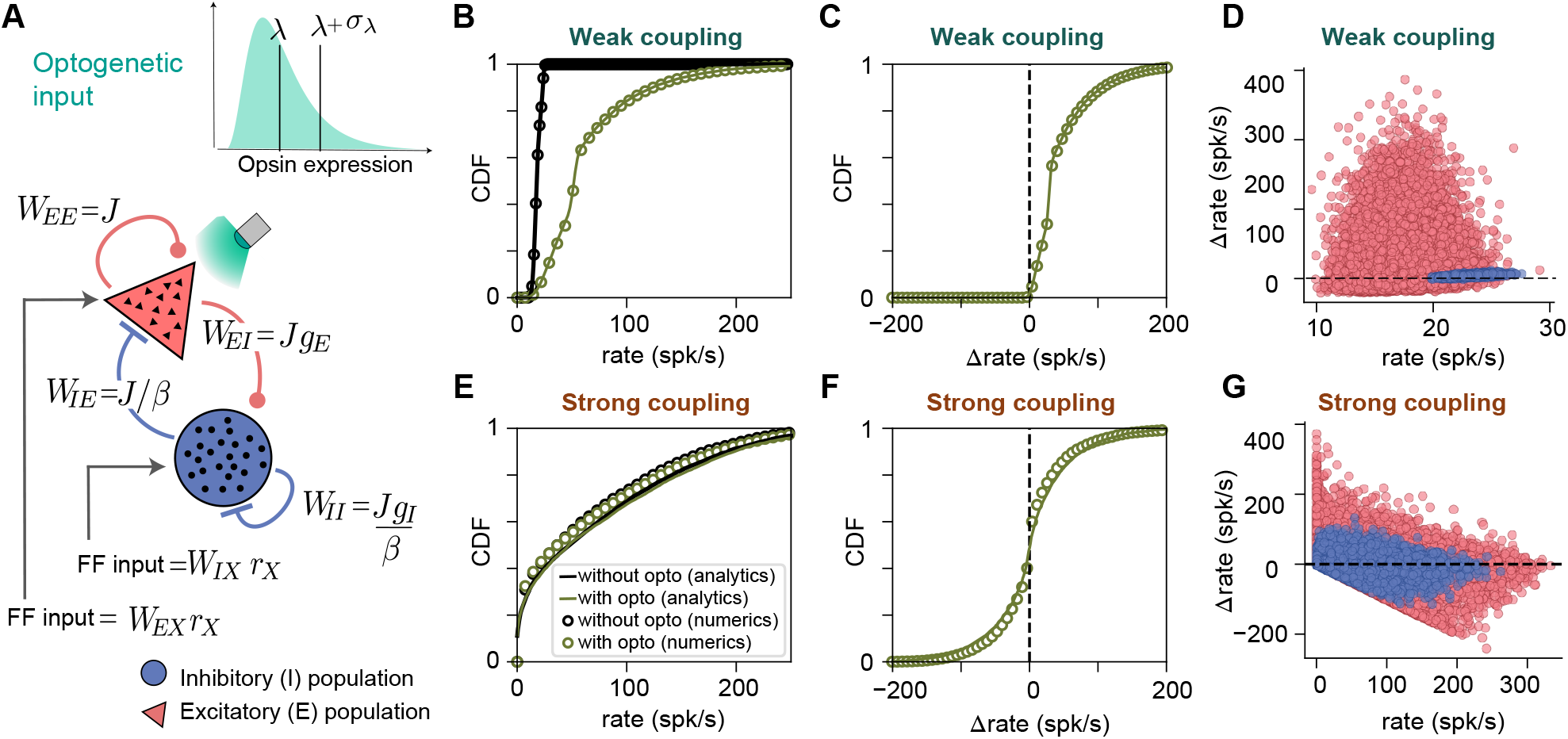
Strong coupling leads to large fractions of suppressed cells, negative covariance and activity reshuffling in randomly connected E-I network models. (**A**) Schematic of unstructured network model, with randomly connected E and I rate units. The network is driven by an external (visual) input, targeting both cell types, and by an optogenetic input targeting only E neurons. Visual inputs follow a Gaussian distribution, while opsin expression in E cells is distributed according to a lognormal distribution. (**B-D**) Responses to optogenetic stimulation of E cells in a weakly coupled network model. In all panels, dots represent numerical simulations while lines are mean field theory predictions. For weak coupling, optogenetic stimulation strongly affects the distribution of firing rates (**B**), with only a weak fraction of suppressed cells (**C**). Changes in firing rates are positively correlated with baseline activity (**D**, Pearson correlation coefficients are 0.1 and 0.6 for E and I cells, respectively). (**E-G**) As in panels **B-D**, but in a strongly coupled network model. For strong enough coupling, optogenetic stimulation does not significantly affect the distribution of rates (**E**). Firing rate changes are broadly distributed, with a mean around zero and a large fraction of suppressed cells (**F**). Firing rate changes are anticorrelated with baseline activity (**G**, Pearson correlation coefficients are −0.3 and −0.2 for E and I cells, respectively). Parameters: *J* = 0.01mV and 0.3mV for weak and strong coupling, respectively; for other parameters see table 1, *Methods*. In all panels, only cells whose response was higher than 1 spk/s in at least one of the two conditions are shown (100% for weak coupling, 43% for strong coupling.) Here and in all modeling figures, unless otherwise specified, results are shown for all cells, both E and I.

We now turn to a deeper investigation of the circuit mechanisms underlying these different aspects of the response of the network to optogenetic stimulation, using mean-field analysis.

### Mechanisms shaping the response to optogenetic inputs in network models

We performed numerical and mean field analyses of the network (see details in *Methods*) to elucidate the circuit mechanisms underlying the simulation results highlighted in Fig. 4. These analyses show that, with strong recurrent connectivity, optogenetic input evokes only a weak change in average firing rate, provided the network is inhibition-dominated (Fig. 5). In particular, numerical simulations and mean field theory show that the average change in firing rate due to optogenetic stimulation decreases when (Fig. 5A-C, first row) the firing rate *r_X_* of the neurons providing external inputs to the network increases, synaptic efficacies *J* become stronger, the strength of inhibition onto E neurons *g_E_* becomes stronger, and/or the strength of inhibition onto I neurons *g_I_* becomes weaker. These results can be understood analytically from the implicit equation defining the mean rates (Eq. (12), *Methods*). We linearize the equation about the fixed point, thus assuming optogenetically induced rate changes in E and I firing rates, Δ*r_E_* and Δ*r_I_*, are small. In the limit of small mean optogenetic input *λ*, and neglecting the contributions coming from quenched disorder in the connectivity and in the opsin expression, we can solve the equations to find

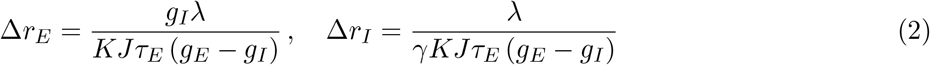

where *K* and *γK* are the average number of E and I recurrent connections per neuron, while *τ_A_* is the membrane time constant of population *A* ∈ [*E, I*]. Eq. (2) captures qualitatively the dependency on *J* and *g_E,I_* observed in simulations (Fig. 5A-C, first row).

**Figure 5:**
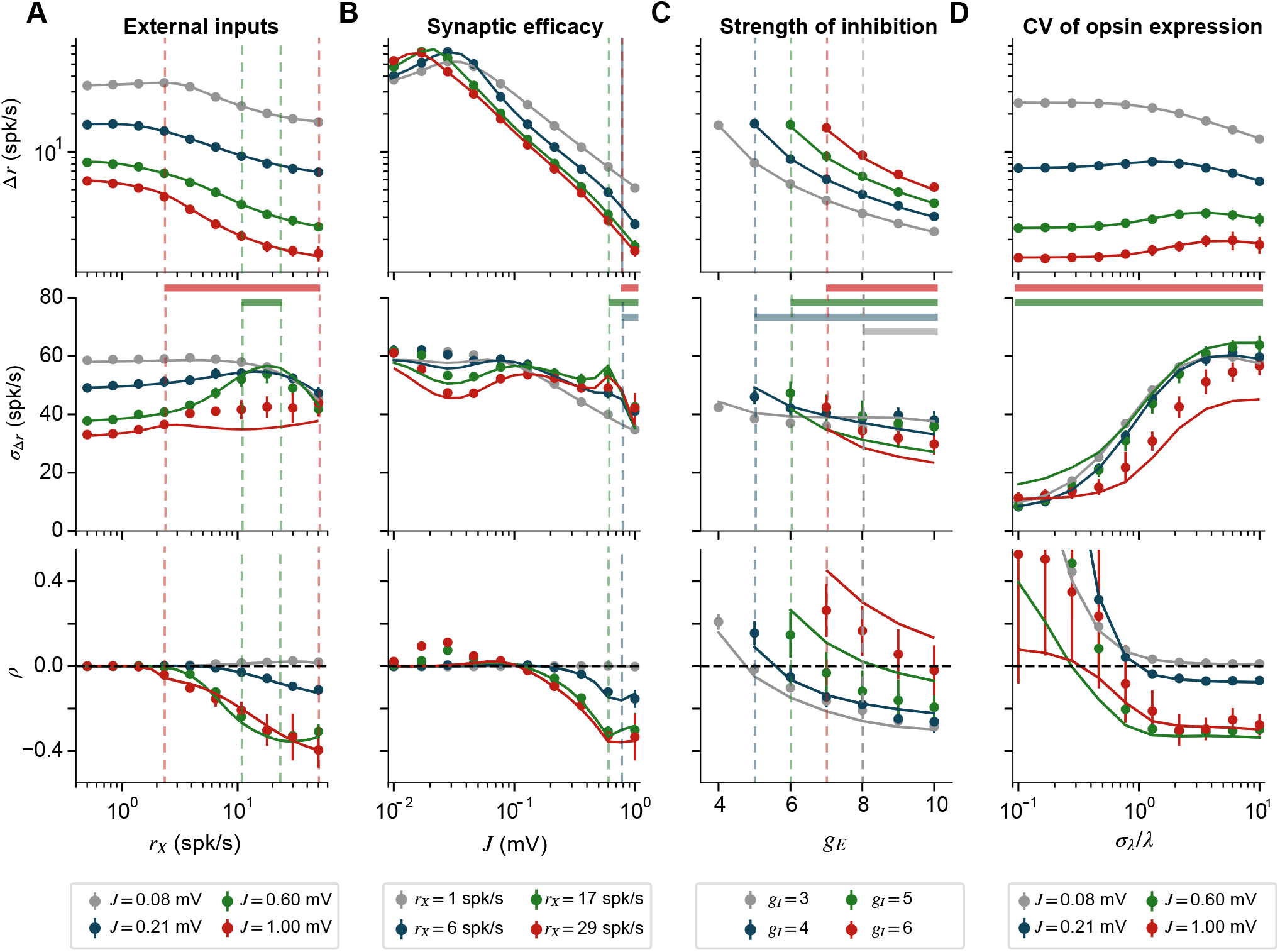
Dependence of the statistics of network response to optogenetic input on network parameters and optogenetic input statistics. (**A**) Statistics of network response to optogenetic stimuli as a function of external input *r_X_*, for different values of recurrent synaptic efficacies (J, colors; *J* scales all weights, ({E,I,X}-to-{E,I}) where X is external input. Panels show (top to bottom): mean change in firing rate Δ*r*; standard deviation of the change in firing rates *σ*_Δ*r*_; normalized covariance *ρ* between baseline rate and change in rates (Eq. 1). Colored bars under the first row indicate, for *J* indicated by color, values of *r_X_* leading to chaotic dynamics. Filled circles with errorbars are mean±sem computed over 20 realizations of the network model. Continuous lines are mean field theory predictions. Increasing *r_X_* generates smaller mean rate change and more negative covariance (except for the smallest J), while only mildly affecting the standard deviation of rate changes. Note that negative covariances can appear both in chaotic and non-chaotic networks (red and green curves, respectively, in lower panel). (**B**) Same as in **A** but as a function of *J*, for different values of *r_X_* (colors). Increasing *J* significantly decreases mean rate change and response covariance, while affecting only mildly the standard deviation of rate changes. (**C**) Same as in **A** but as a function of the strength of inhibition to E cells *g_E_*, for different values of the strength of inhibition to I cells *g_I_*. Stronger inhibition to E cells and weaker inhibition to I cells both lead to lower mean rate changes and more negative covariance of rate change with baseline rate, while only mildly affecting the standard deviation of rate changes. Networks with *g_E_* < *g_I_* are not analyzed because they are either unstable or multistable (Sanzeni et al., 2020). (**D**) Same as in **A** but as a function of the coefficient of variation (CV) of the distribution of opsin expression *σ_λ_/λ*, for different values of *J*. The CV weakly affects the mean rate changes, but strongly affects the standard deviation of the rate changes and the response covariance. In particular, heterogeneous opsin expression (*σ_λ_/λ* > 1) is required to generate broad distributions of responses and negative covariance. Simulation parameters are: *σ_λ_/λ* = 2 in panels **A-C**; *g_E_, g_I_* = 8, 3 in panels **A,B,D**; *r_X_* = 10spk/s and 18spk/s in panel **C** and **D**; *J* = 0.6mV in panel **C**.

Strong recurrent input can largely cancel the mean optogenetic input, but cannot cancel the cell-to-cell fluctuations in input induced by opsin expression heterogeneity (Renart et al., 2010; van Vreeswijk and Sompolinsky, 1998). This produces high response heterogeneity (Fig. 5, second row) that increases with opsin expression variability (*σ_λ_*, Fig. 5D, second row), and only weakly depends on coupling strength (Fig. 5A–C, second row). Heterogeneous opsin expression and strong coupling also lead to a large number of suppressed cells (Figs. 4.)

The combination of strong coupling and a broad distribution of opsin expression naturally leads to a normalized covariance *ρ* closer to −0.5 (Fig. 5, third row) and to network reshuffling (Fig. 4E–G). The fact that *ρ* becomes negative is due to an interplay between single neuron nonlinearities and the strong inhibitory feedback canceling the mean network response to optogenetic input. Strong couplings leads to a broad distribution of baseline rates and therefore a large fraction of cells with very low baseline rates. Upon heterogeneous optogenetic stimulation, the non-negative nature of firing rates biases the response of cells with very low baseline rates toward excitation (see Fig. 4G). To compensate for this bias, and generate a close to zero mean change in rate, the circuit produces, on average, suppression in neurons with higher baseline activity (see Fig. 4G). This leads to a negative correlation between baseline activity and response to optogenetic input, i.e. a negative ρ. This negative correlation leads in turn to a negative correlation between baseline synaptic inputs and changes in inputs due to optogenetic drive (see Eq. (10).

Previous theoretical works have shown that the dynamics of rate networks becomes chaotic when recurrent connections are strong (Harish and Hansel, 2015; Kadmon and Sompolinsky, 2015; Sompolinsky et al., 1988). Consistent with these results, we observed the emergence of chaotic dynamics in our simulations as *J* and/or *r_X_* increase (Fig. 5, lines under the first row). Note that in our simulations chaos is heterogeneous across cells, with a broad distribution of temporally-averaged firing rates that reflects the quenched disorder in the inputs and synaptic connectivity. Since negative covariance and neuronal reshuffling emerge at large contrast, we wondered if the appearance of these features is related to chaotic dynamics. To answer this question, we have systematically compared the regions of parameter space in which negative covariance and chaotic dynamics appear (supplementary information Fig. S2) but found that they only partially overlap. These results show that negative covariance and chaotic dynamics are two distinct phenomena that both emerge in networks of strongly coupled neurons.

### Randomly connected network models capture experimental recordings in mice and monkeys

Optogenetic responses in mice and monkeys have qualitatively similar features, which we summarized above with the term “network reshuffling”, but also important quantitative differences (Figs. 1 and 2). Compared to monkeys, optogenetic responses in mice had a larger mean over standard deviation ratio and weaker normalized covariance. However, when these quantities are plotted as a function of mean baseline population rate (in the absence of optogenetic stimuli), these differences seem to largely arise because of different ranges of baseline firing rates, possibly due to a difference in visual input strengths in the two species (Fig. 3A–B). Thus, the strength of external inputs might be the main factor explaining differences between mice and monkeys. To test this hypothesis, we investigated if a single unstructured network model, with only the strengths of external stimuli differing between the two species, can capture the responses observed in each.

We fitted our network model to recordings in mice and monkeys assuming common network parameters (see methods), but allowing for different external (*r_X_*) and optogenetic (*L*) inputs. This model captures the main features of the recordings (Fig. 6A, dashed lines): weak mean optogenetic response; large standard deviation of optogenetic response; and increasingly negative normalized covariance with contrast of the visual stimulus. It also captures key differences between mice and monkeys: a lower visual response, and a larger ratio between mean and standard deviation of the optogenetic response in mice; and a more negative normalized covariance in monkeys. These results support the idea that the strength of external inputs is a key factor shaping differences between the two species.

**Figure 6:**
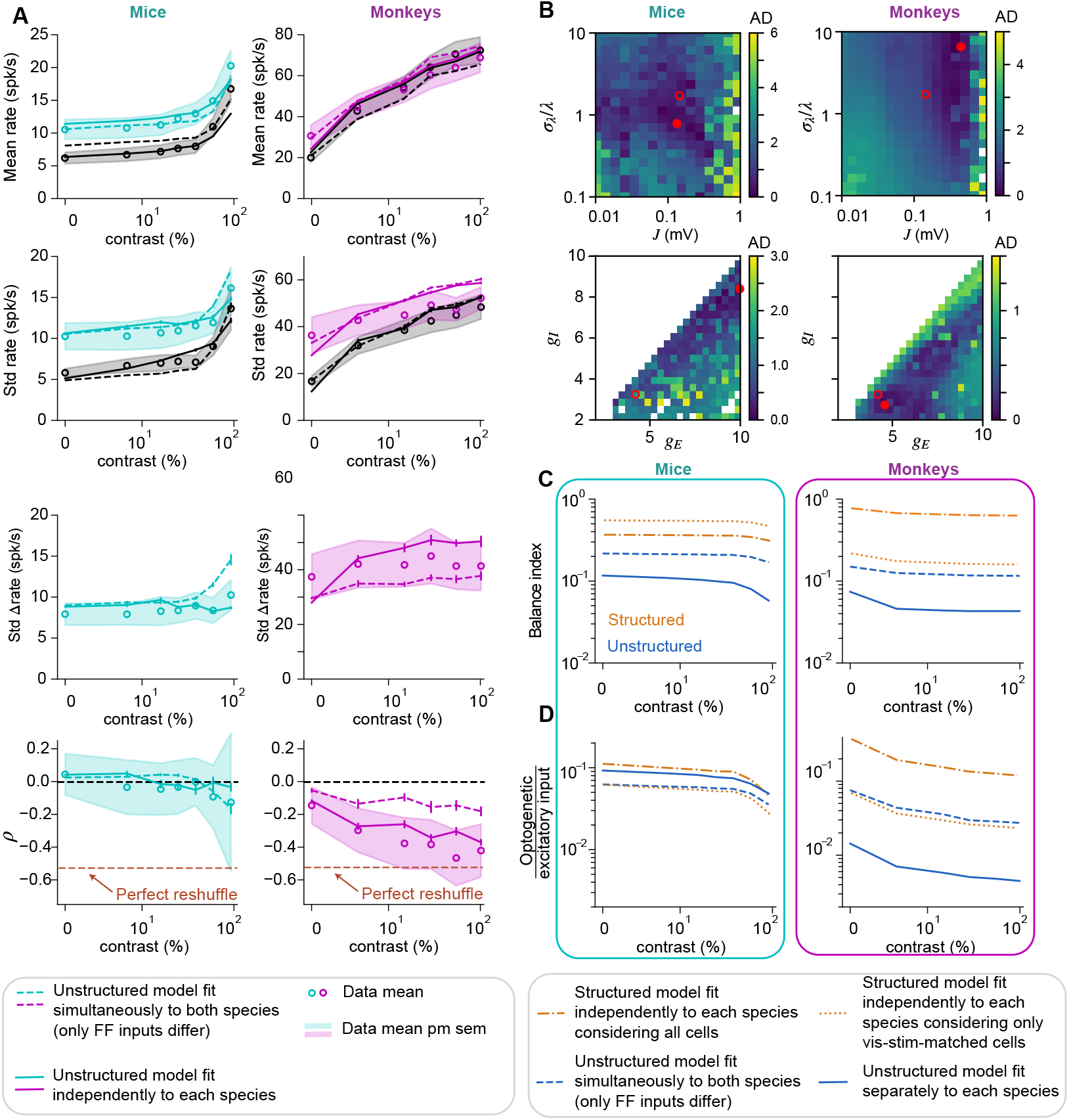
Fitting mice and monkey data with the network model. (**A**) Comparison between data (circles represent means, shaded regions are mean±sem) and network model with parameters that best fit data (dashed and continuous lines) in mice (left) and monkeys (right). Dashed lines represent best fits obtained with the same network model in mice and monkeys, but different external and optogenetic inputs. Continuous lines represent best fits obtained with different network models. Model plots are obtained from simulations, averaging over 20 realizations. Network models reproduce key features of experimental data, including: weak mean changes in firing rate; broad distribution of responses; and increasingly negative covariance with contrast of the visual stimulus. (**B**) Average discrepancy (AD, measured as log_10_ of the ratio between the actual and the minimum error) between best model predictions and experimental data for mice (left) and monkeys (right) obtained with models where two parameters are constrained to the values specified by both axis. Red circles indicate optimal parameters, used to generate panels **A** (open circles: same parameters in mice and monkeys; filled circles: different parameters in mice and monkeys). In both species, consistent with Fig. 5D, better fits are obtained for *σ_λ_/λ* > 1, i.e. highly heterogeneous optogenetic expression, and for biologically realistic values of J, in the range of [0.1 – 1]mV. (**C**) Inferred values of the balance index (Ahmadian and Miller, 2021), i.e. the ratio between net input current (excitatory minus inhibitory) and the total excitatory input in response to visual stimuli. Fits in structured models (orange lines) are less strongly coupled that fits in the unstructured model. (**D**) Inferred values of the ratio between optogenetic input and the total visually-driven excitatory input. Both in mice and monkeys, the unstructured model (blue lines) requires a small mean optogenetic input, relative to visually-driven excitatory input, to capture experimental data. In structured models, data can be captured with a larger optogenetic to excitatory input ratio.

At the same time, the model fails to capture quantitatively the mean optogenetic response and the amplitude of the negative normalized covariance in monkeys. These discrepancies suggest that differences in the network structure or the visual or optogenetic input distributions might be important to fully capture the observed response properties. To test this, we repeated the fitting procedure in unstructured models, allowing different connectivity and input parameters for mice and monkeys, and found that this model gives a better description of experimental recordings, including the strongly negative covariance seen in monkeys for high contrast (Fig. 6A, continuous lines; relative error decreases averaged over the quantities plotted in Fig. 6A were 22% in mice and 5% in monkeys). The best fit parameters are characterized by weaker overall coupling *J*, weaker inhibition (weaker overall *I* →E strength, *J_g_E__*, and weaker ratio of I→E to I→I strength, *g_E_/g_I_*), and less heterogeneous optogenetic expression (smaller *σ_λ_/λ*) in mice than in monkeys.

Previous theoretical work (Ahmadian and Miller, 2021; Rubin et al., 2015; Sanzeni et al., 2020) has suggested that cortical circuits operate in a loosely balanced regime, characterized by the net input to cells (excitatory minus inhibitory) being comparable to its individual (excitatory or inhibitory) components. This can be quantified by the ratio of the two – net input over excitatory input – which is called the balance index (Ahmadian and Miller, 2021). In our models, using the parameters inferred from data, we found a smaller balance index in monkeys than in mice (Fig. 6C). Recordings in mice were consistent with the underlying network being loosely balanced at low and intermediate contrast (balance index greater than 0.1). For the model in which mice and monkeys were jointly fit, somewhat tighter balance was observed at high contrast in mice and for all contrasts in monkeys (Fig. 6C, dashed line), while monkeys showed very tight balance in the model in which they were independently fit (Figure 6C, filled line).

The unstructured model also requires the ratio of mean optogenetic input to visually evoked excitatory input to be small (~ 0.1), and to be very small (≲ 0.01) for the independent fit to monkeys (Figure 6D). The latter is related to tight balancing. The small balance index needed for the independent fit to monkeys indicates that the mean inhibitory recurrent input cancels most of the mean excitatory input (*e.g*. a balance index of 0.04, Figure 6C, means that all but 1/25 of the mean excitatory input is cancelled). In particular, both the visual and the optogenetic input are strongly cancelled, by similar factors. For the mean optogenetic response to be reduced by cancellation to near zero, while the mean visual response remains finite, the mean optogenetic input must be much smaller than the mean visual input. Thus, by both measures – the balance index, and the relative mean size of optogenetic vs. visual input – the nearly perfect reshuffling seen in monkeys at higher contrast (very small Δ*r* and strongly negative normalized covariance *ρ*) appears to require tight balancing, which in turn means that coupling is very strong (Figure 5A,B; Ahmadian and Miller, 2021.

### Structured connectivity leads to reshuffling with weaker coupling

The unstructured models studied to this point ignored the difference between tuned visual input, which specifically targets a set of cells with similar feature and spatial selectivity, and untuned optogenetic input, which targets cells in a given cortical region regardless of their selectivity. It similarly ignored the tuning of connectivity, which decreases with cortical distance (Angelucci et al., 2002; Rossi et al., 2020) and, at least in mice, with difference in feature preference (Cossell et al., 2015; Ko et al., 2011). We reasoned that a tuned stimulus would excite cells with matched tuning, which might preferentially excite one another and/or inhibit others; whereas an untuned stimulus would tend to excite and inhibit all cells equally, which could give a very small or even negative response. With such a mechanism, the network might not need such tight balance or small optogenetic input to explain reshuffling.

To test these ideas, we constructed alternative, structured models. We assumed that neuronal selectivity was determined by a single feature, preferred orientation. Connection strengths decayed as a product of Gaussian functions of the physical distance between neurons and of the difference in their preferred orientations. We used a spatially salt and pepper organization of preferred orientation in the mouse and a quasi-periodic map in the monkey, as reported experimentally (Hubel and Wiesel, 1977; Kaschube et al., 2010; Ohki et al., 2005, 2006). Visual input was also a product of Gaussian functions of the distance between receptive field center and stimulus location and of the difference between preferred orientation and stimulus orientation.

We fit these structured models to the data and found that the same quality of fit could be obtained as with the unstructured models fit independently to mice and monkeys (see Fig. S3). However, this fit for monkeys is obtained with a much larger balance index than the independently fit unstructured models: ~ 5 times larger if the index is computed only over cells whose preferred orientation matches that of the stimulus, (*visual-stimulus-matched* cells; Fig. 6C, orange dotted lines, see also Fig. S4) and even larger if computed over all cells (orange dash-dotted lines). Similarly, the structured models can match the monkey’s nearly perfect reshuffling with a much larger ratio of mean optogenetic to visual input than the unstructured models (~ 10 times larger for visual-stimulus-matched cells, even larger for all cells; Fig. 6D, see also Fig. S4). As in the unstructured model (Fig. 4), the structured model requires sufficiently strong coupling for the mean optogenetic response to be suppressed (Fig. S5), although less strong than is required in the unstructured model (Fig. S4). In sum, the structured models can produce strong reshuffling with significantly weaker balance and stronger optogenetic input than the unstructured models.

## Discussion

### Summary of the results

We analyzed electrophysiological recordings of mice and monkey V1 while the animals viewed visual stimuli, with or without concurrent optogenetic stimulation of E neurons. In both species, and for all contrasts of visual stimuli, optogenetic stimuli generate a broad distribution of responses, whose mean is considerably smaller than the standard deviation. This large heterogeneity of optogenetic responses results, in particular, in large fractions of suppressed cells. The main quantitative differences between the two species are that, in monkeys, the firing-rate responses to visual and optogenetic stimulation, and the fraction of suppressed cells, are larger, while the mean optogenetic response is smaller. Despite this strong optogenetic modulation of the activity of single neurons, the distribution of rates across the population is unchanged by optogenetic stimulation in monkeys and, for high visual contrast, in mice. This “reshuffling” of rates is specific to optogenetic stimuli; visual stimuli shift the distribution toward higher rates. Intriguingly, statistics of optogenetic responses across the two species form a single, continuous function of baseline (visually-induced) population activity, with mouse and monkey responses corresponding to lower and higher baseline activities, respectively.

Why does optogenetic stimulation drive large single-cell responses but little or no mean response? To address this, we first studied an unstructured, randomly connected network model of E and I neurons. We found that the “balancing” that occurs in sufficiently strongly coupled E/I networks (Ahmadian and Miller, 2021; van Vreeswijk and Sompolinsky, 1998) – the dynamic cancellation of external input by recurrent input – can produce this effect. Balancing suppresses the mean input, but not the variation in input across cells. Accordingly, when the heterogeneity of optogenetic input (i.e. of opsin expression) is comparable to or larger than the mean optogenetic input, the mean response is suppressed while individual cells show large responses. In this model, optogenetic input is distinguished from visual input in (1) targeting E, rather than both E and I, cells, (2) having a much smaller mean, and (3) having larger variability relative to mean. The small relative size of mean optogenetic input is a possible limitation of the model (further discussed below), and is required to have a substantial mean visual response with nearly zero mean optogenetic response, despite both being suppressed by balancing. Finally, for strong reshuffling – invariant firing rate distributions despite strong, heterogeneous single-cell responses – there must be a strong negative covariance between optogenetic and baseline responses. This arises because neurons with low baseline rates are biased by rectification to positive optogenetic responses, and the circuit’s dynamic suppression of the mean response therefore causes suppression of neurons with higher baseline activity. With appropriate adjustment of parameters, this simple model could be well fit to the mouse and the monkey data. A second possible limitation of this model is that, to achieve strong reshuffling, the network balance of excitation and inhibition must be tighter than has been estimated from experiments (Ahmadian and Miller, 2021), also discussed below.

We then incorporated the fact that the visual input is tuned, *e.g*. it targets cells with preferred orientation near the stimulus orientation, while the optogenetic input targets all E cells. We reasoned that a tuned input might lead to stronger excitation and/or weaker inhibition than an untuned input (e.g. as in the model circuit of Troyer et al., 1998), allowing suppression of the mean response under less stringent conditions. Studying such a model with structured connectivity (feature- and distance-dependent), we found that it could achieve equally good fits to the data as the unstructured model, but with looser balancing in the reshuffled cells, and with larger optogenetic input relative to visual.

### Theoretical implications: Coupling strength in cortex

Balanced network models (Ahmadian and Miller, 2021; Ahmadian et al., 2013; Amit and Brunel, 1997; Brunel, 2000; Renart et al., 2010; Roxin et al., 2011; Rubin et al., 2015; Sanzeni et al., 2020, 2022; van Vreeswijk and Sompolinsky, 1996, 1998) have been successful in reproducing several ubiquitous features of cortical activity, including highly irregular firing (Compte et al., 2003; Softky and Koch, 1993), excitation/inhibition balance (Haider et al., 2006), and broad distributions of firing rates (Buzsaki and Mizuseki, 2014; Hromádka et al., 2008; O’Connor et al., 2010). These properties can arise either in networks that are tightly balanced or loosely balanced (Ahmadian and Miller, 2021), meaning that the net input after cancellation of excitation and inhibition is respectively much smaller than, or comparable in size to, the cancelling factors (balance index roughly ≲ 0.1 or ≳ 0.1, respectively). Tighter balance corresponds to stronger coupling. Here, we showed that such networks can reproduce the surprising effect of strong reshuffling. This provides further evidence that recurrent connectivity in V1 is powerful, and stabilized by strong inhibition. A key property of the loosely balanced regime, as exemplified by the supralinear stabilized network (SSN) model (Ahmadian and Miller, 2021; Ahmadian et al., 2013; Rubin et al., 2015), is that network behavior depends on the network’s activation level. In particular, in the region over which the neuronal input/output function is supralinear, increasing activation yields increasingly strong coupling. In our models, this accounts for the strong dependence of the results of optogenetic stimulation on the state of the network prior to stimulation, which in the experiments described here is controlled by the contrast of the visual stimulus.

The unstructured model illustrates basic mechanisms, but requires relatively tight balance (balance index (b.i.) of best fits at high contrast: monkey, ~ 0.03; mouse, ~ 0.05). The structured model shows similar mechanisms can operate with looser balance (b.i. of reshuffled cells in best fits, high contrast: monkey or mouse, ~ 0.2), consistent with estimates from experimental data of loose balance (Ahmadian and Miller, 2021). Note, however, that those estimates were based on cat and mouse V1; it is possible that monkey V1 has tighter balance, *e.g*. due to its higher firing rates (although extracellular recording may overestimate firing rates, Barth and Poulet, 2012).

An interesting feature of our analysis is that reshuffling and chaos often coexist (Engelken et al., 2020; Harish and Hansel, 2015; Kadmon and Sompolinsky, 2015; Sompolinsky et al., 1988). This makes sense since both require strong coupling. We note however that our results do not provide evidence of chaos in cortical circuits, since non-chaotic networks reproduce all the main features in the data.

### Mice vs monkeys

Our work provides to our knowledge the first systematic comparison of how optogenetic stimuli modulate visual responses in mice and monkeys. It shows that baseline firing plays a key role in determining the statistics of neural responses, and suggests that the different activity levels, due to different strengths of external input, could underlie differences observed in the two species. When key statistics were plotted as a function of baseline rate, curves traced by mice and monkey data seem to lie on the same underlying curve, with mice occupying the low rate region while monkeys occupy the high rate region. Consistent with this observation, fitting data from both species with a network model with the same parameters, but different input strengths, gives a reasonable description of the data, though the quality of the fit improves when network parameters are allowed to differ between the two species.

A recent electron microscopy study performed a detailed comparison of reconstructed neurons and their synaptic connections in mouse and primate V1 (Wildenberg et al., 2021). Surprisingly, synaptic connectivity in primates was found to be much more sparse than in mice, with a larger ratio of excitatory to inhibitory synapses in mice. The larger ratio of excitatory to inhibitory synapses in mice seems consistent with the weaker inhibition we found in mice. However, the weaker anatomical coupling in monkeys than in mice seems at first sight at odds with our conclusions that monkeys appear to be in a more strongly coupled regime than mice. It is possible that the physiological strengths of the connections are stronger in monkeys, sufficiently so that monkey connections are equal or larger in overall strength to those in mice. In addition, strong coupling can arise from stronger activation levels as well as from stronger synaptic connections, and the former are larger in monkeys. Furthermore, both the unstructured and the structured models reproduce well the data in broad regions of parameter space (see, in Fig. 6B, the broad blue regions which indicate reasonable fits in a broad range of values of J, which scales both synaptic strengths and input strengths).

Running strongly enhances visual responses in mouse V1 (Niell and Stryker, 2010). Recent experiments found that, in V1 of a species of monkey (the marmoset), running does not increase the mean response to visual stimuli, but strongly modulates single-neuron responses, with a large fraction of cells suppressed (Liska et al., 2022). This is similar to the difference we have found in mice vs. monkeys in responses to optogenetic stimuli (although the mouse mean response increase for running is considerably stronger), suggesting possible common mechanisms: the weaker running modulation in monkeys than in mice could, at least in part, be a consequence of stronger coupling in monkeys.

### Relations with previous experimental works and computational consequences

The response properties of cortical networks to optogenetic inputs we have found here might also be relevant to understanding responses of neurons observed in other conditions. Large numbers of suppressed cells in response to optogenetic activation of pyramidal cells have been reported in experiments using one-photon stimulation in mouse visual cortex (O’Rawe et al., 2022), ferret visual cortex (Wang et al., 2022), and targeted two-photon stimulation of small ensembles in mouse barrel cortex (Dalgleish et al., 2020). Weak changes in mean population activity together with large changes in individual neuronal firing rates also characterize the response of cortex to some modulatory inputs (Sherman and Guillery, 1998), *e.g*. in behavioral modulations of activity by head movements (Bouvier et al., 2020) and running (Liska et al., 2022). Similar changes are seen after learning in experiments involving brain-machine interfaces (Engelhard et al., 2019).

Some recent works have reported significant changes in mean firing rate with optogenetic stimulation of E cells (Andrei et al., 2019; Chen et al., 2022) in monkeys. Many possible factors might contribute to differences from our results, including strength, timing, specificity, and spatial spread of optogenetic stimulation, visual stimulus strengths, match or lack thereof between the stimulus and neuronal preferences, recording methods, and behavioral paradigms used. In particular, Andrei et al. (2019) used very weak stimulus contrast, for which both our data and our models indicate weaker suppression, and used orientation-specific optogenetic stimulation which, like tuned visual input, may be able to drive mean activity. They could roughly match visual stimulus orientation, but not other stimulus features, to neuronal preferences. They also analyzed only cells with significant laser response, while we included all recorded cells in our analysis. Chen et al. (2022) used wide-field imaging to record the pooled activity of a large population of neurons with diverse tuning properties. Surprisingly, they generally found no mean visual response given the optogenetic stimulation, which might suggest their mean optogenetic input was very large compared to their mean visual input, the opposite of what we have concluded about our monkey experiments. Both experiments were done in the context of behavioral (detection or discrimination) tasks. Training mice to detect optogenetic stimuli can dramatically increase network responses to those stimuli (Akitake et al., 2022); it is conceivable that behavioral training could alter network responses in important ways.

### Limitations and future directions

For the best fit match to the monkey at high contrast, the unstructured model had a ratio of optogenetic to excitatory visual input of about 5 × 10^-3^. The structured model relaxed this requirement, but its ratio for visual-stimulus-matched cells was still small, about 2 × 10^-2^. Are these plausible? In layer 4 of mouse V1, peak visually-evoked excitatory current was estimated to be 60-150pA across cells (Li et al., 2013; Lien and Scanziani, 2013). In cultured neurons, optogenetically-evoked currents are 100’s to 1000’s of pA (Mattis et al., 2011), but light is severely attenuated with transmission through cortex, by 50% over about 40 microns (Yona et al., 2016). Thus, it is difficult to assay whether the model’s requirement is met *in vivo* in cortical cells, but this could be a problem for the present scenarios.

Though the current structured models succeed in explaining the experimental results with looser balance and stronger relative optogenetic input, we lack a detailed understanding of the exact mechanisms by which network structure affects reshuffling. It also remains uncertain whether different mechanisms or requirements might be revealed by considering other forms of structure, such as neurons with different preferred spatial phases (*e.g*. Kayser and Miller, 2002; Troyer et al., 1998) or preferred directions (Rossi et al., 2020). Future work will examine these questions.

### Conclusions

The over 70 million years of evolution separating mice and monkeys (Siepel, 2009) have produced substantial differences in the structure of their cerebral cortices, e.g. in the presence of orientation and ocular dominance columns in primates (Hubel and Wiesel, 1977) but their absence in rodents (Dràger, 1974; Ohki et al., 2005; Van Hooser et al., 2005) the size of their receptive fields (Van Den Bergh et al., 2010), and their connection statistics (Wildenberg et al., 2021). Our analysis of optogenetic modulation of visual response in V1 shows that, despite these differences, mice and monkeys may share a common remarkable resistance of their cortex to optogenetic inputs at the population level, in spite of large changes in single neuron firing rates. At the same time, we find a key difference may be that monkeys are in a more strongly coupled regime of higher firing rates, with correspondingly stronger resistance. These results might more generally yield insight into circuit mechanisms underlying differences in the effects of sensory or driving vs. modulatory inputs on cortical responses.

## Methods

### Experimental methods

#### Mice

A detailed description of the experimental methods is given in (Histed, 2018). In brief, neurophysiological data from Emx1-Cre animals (*n* = 4) were collected. Animals kept on a monitored water schedule were given small drops of water (~1 μl) every 60–120 s during recording to keep them awake and alert. The visual stimulus, a Gabor patch with spatial frequency 0.1 cycle/deg and sigma 12.5 deg, were presented for 115 ms [full width at half maximum (FWHM) intensity], and successive visual stimuli were presented every 1 s. Optogenetic light pulses were delivered on alternating sets of 10 stimulus presentations (light onset 500 ms before first stimulus; offset 500 ms after end of last stimulus; total light pulse duration 10.2 s). A 1-s delay was added after each set of 10 stimulus presentations. Extracellular probes were 32-site silicon electrodes (Neuronexus, probe model A4×8). Channelrhodpsin-2 (ChR2) was expressed in E neurons (as described in Histed and Maunsell, 2014) using viral (AAV-EF1a-DIO-ChR2-mCherry, serotype 2 or 8; http://openoptogenetics.org) injections into the Emx1-Cre (Gorski et al., 2002; #5628, Jackson Laboratory) line. Virus (0.25–1.0 μl) was injected into a cortical site whose retinotopic location was identified by imaging autofluorescence responses to small visual stimuli. Light powers used for optogenetic stimulation were 500 μW/mm2 on the first recording session; in later sessions, dural thickening was visible and changes in firing rate were smaller, so power was increased (maximum 3 mW/mm2) to give mean spontaneous rate increases of approximately ~5 spk/s in that recording session. Optogenetic light spot diameter was 400–700 *μ*m (FWHM) as measured by imaging the delivered light on the cortical surface.

Spike waveforms were sorted after the experiment using OfflineSorter (Plexon, Inc.). Single units (SU) were identified as waveform clusters that showed clear and stable separation from noise and other clusters, unimodal width distributions, and interspike interval histograms consistent with cortical neuron absolute and relative refractory periods. To compute neurons’ visual responses, we counted spikes over a 175-ms period beginning 25 ms after stimulus onset. Zero-contrast responses were computed from the 175 ms period immediately preceding stimulus onset.

#### Monkeys

A detailed description of the experimental methods is given in (Nassi et al., 2015). In brief, two adult male rhesus macaques (Macaca mulatta) were each implanted with a custom titanium head post and silicone-based artificial dura recording chamber over V1. We injected a VSVg-pseudotyped lentivirus carrying the C1V1-EYFP gene behind the 1.3 kb CaMKII*α* promoter (lenti-CaMKII*α*-C1V1E162T-ts-EYFP; titer 3 × 1010 TU/ml) into a single location in V1 in each of the two monkeys (monkeys A and M) while they were anesthetized and secured in a stereotactic frame. Injections of the same viral construct were made into V1 of one additional monkey in order to assess specificity of viral expression to E neurons. In a second, distant location in V1 of monkey A, we injected an adeno-associated virus carrying the eArch3.0-EYFP gene behind the 1.3 kb CamKII*α* promoter (AAV5-CamKII*α*-eArch3.0-ts-EYFP; titer 4 × 1012 VP/ml).

Monkeys were alert and head restrained during all experiments. Single- and multi-unit waveforms were recorded using thin glass-coated tungsten electrodes (Alpha Omega, Engineering Inc., Nazareth, Israel). After isolating a single-unit or multi-unit cluster, we first assessed sensitivity to optogenetic stimulation. We randomly interleaved different stimulation intensities. Stimulation on each trial was continuous and lasted for 200 ms. Each condition was repeated at least five times. For a subset of light-sensitive units, we proceeded to measure responses to simultaneous optogenetic stimulation and visual stimulus presentation. The presented visual stimuli consisted of circular patches of drifting sinusoidal gratings whose size was matched to each recorded unit’s preferred size (median diameter = 0.63deg), of mean luminance matching the surround (42 cd-m2) at the optimal orientation and spatial and temporal frequencies. Each stimulus condition was presented at least five times. For simultaneous optogenetic stimulation and visual stimulus presentation, we varied the contrast of the presented gratings in log steps (0%, 6%, 12%, 25%, 50%, and 99%) and stimulated with four different intensities including zero. Visual contrast and stimulation intensity were randomly interleaved. Optogenetic stimulation onset occurred simultaneously with the onset of the visual stimulus and lasted 200ms. The visual stimulus was always presented for at least 200ms, but in some cases lasted 300ms. Response was measured as the mean response over the interval 100-200ms after stimulus onset. Here, we show results only for two optogenetic intensities, zero and the largest intensity, but similar results hold also for intermediate intensities.

### Mathematical Methods

#### Network Models

For all models, the single neuron response function (firing rate vs input current relationship) was taken to be the f-I curve of leaky integrate-and-fire neurons driven by white noise (Amit and Brunel, 1997; Amit and Tsodyks, 1991; Sanzeni et al., 2020; Siegert, 1951). Specifically, the firing of a neuron belonging to population *A* ∈ [*E, I*] in response to an input *x*, is given by

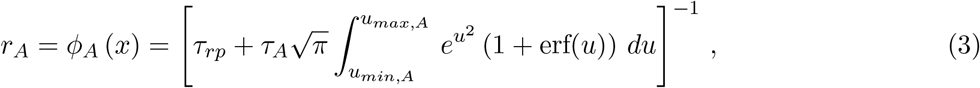

with

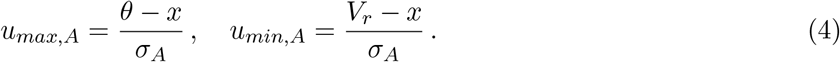

In Eq. 3, *τ_rp_* indicates the single neuron refractory period so that 1/*τ_rp_* is the maximal single neuron firing rate; *τ_A_* is the membrane time constant; *σ_A_* is a parameter controlling the smoothness of the transfer function. Throughout the paper, we assumed the values *τ_rp_* = 2ms, *θ* = 20mV, *V_r_* = 10mV, *τ_E,I_* = 20, 10ms and *σ_E,I_* = 10mV.

Indicating with 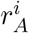 and 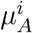 the rate and the total input (recurrent + external) of the *i*-th cell in population *A* ∈ [*E, I*], activity in the network evolves in time according to the equation

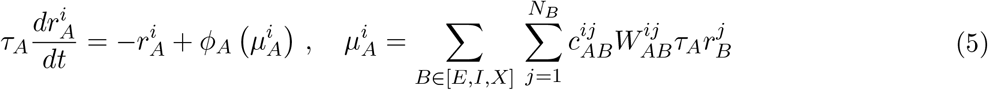

Here 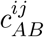 is the adjacency matrix (equal to 1 or 0 according to whether or not there is a connection from cell *j* of type *B* to cell *i* of type *A*) and *N_B_* is the number of neurons in population *B*, where (*X*) indicates the externally driven input, *i.e*. visual input. We always use *N_X_* = *N_E_* = 4 *N_I_*.

Throughout the paper, we assumed uniform *W*’s, 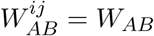, where the weights *W_AB_* are parameterized as follows:

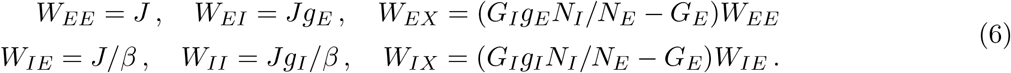

These definitions were chosen so that the parameters *G_I_* and *G_E_* give the gain of the E and I population in the strong coupling limit (see (Sanzeni et al., 2020)).

With optogenetic input, the network dynamics becomes

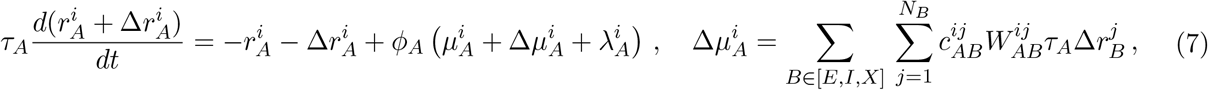

where 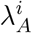 represents the optogenetic input of the *i*-th neuron in the network, which varies according to the opsin expression in that neuron, while 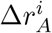 and 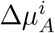 indicate the change in rate and in the recurrent inputs produced by optogenetic stimuli.

#### Unstructured Models

We investigated large randomly connected networks of E and I rate neurons. In the theoretical analysis of the model (mean field theory described below and simulations shown in Figures 4 and 5 and in supplementary information), the network connectivity was assumed to follow an Erdös-Rényi statistics, i.e. each element of the adjacency matrix 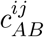 was 1 with probability *p* = 0.1 and 0 otherwise. The mean and variance of the number of pre-synaptic neurons from population B per cell are given by *K_B_* = *pN_B_* and 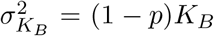, respectively. We define *K* ≡ *K_E_* = *pN_E_*, and use the notation *γ* = *K_I_/K_E_*. In fitting the model to data (Figure 6), we considered more general statistics by generating 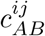 as follows: for each row *i*, a random number, *K_i_* (Gaussian distributed, with mean *K_B_* and standard deviation *σ_K_B__* = *CV_K_* × *K_B_*) of elements were set to 1, with autapses not allowed, and the other elements were set to 0 (where *CV_K_* is a constant that is identical for *B* = *E, I*, or *X*).

#### Structured Models

For the structured models, we place cells onto evenly-spaced discrete locations on a 25 × 25 periodic grid, so that there are *N_G_* = 625 grid points, with 8 E and 2 I cells per grid location. The grid is a square of side dl = 1mm. For the monkey, we introduce a smooth aperiodic orientation map using a method similar to that of Kaschube et al. 2010 by summing 500 plane waves of the form exp[*i*(*k cos*(*jπ*/500)*x* + *k sin*(*jπ*/500)*y* + *φj*)] for *j* = 0, 1,…, 499. The wavenumber *k* is chosen to be 6*π/dl* such that the wavelength is *dl*/3, and the phase *ϕ_j_* is chosen as a random angle for each *j*. The preferred orientation at a point in the grid is calculated as half the phase of the sum of plane waves at that point, such that the orientations are between 0° and 180°. For the mouse, we generate a salt-and-pepper orientation map by randomly shuffling a smooth orientation map generated as described above. We let *θ*(**x**) be the preferred orientation at two-dimensional grid position **x**.

The probability of a connection from a cell at position **y** of type *A, A* ∈ *E, I*, to any cell at position **x** is computed as follows. We let *d*(**x**, **y**) be the shortest distance across the periodic grid between **x** and **y**, and *dθ*(*θ*(**x**), *θ*(**y**)) be the shortest distance around the 180° circle of orientations between *θ*(**x**) and *θ*(**y**). We let *n_A_* be the number of cells of type *A* at a single grid position (*n_E_* = 8, *n_I_* = 2). We then form the functions 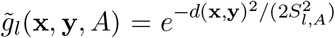 and 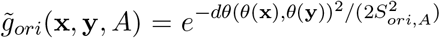, and normalize each by its mean to obtain the product

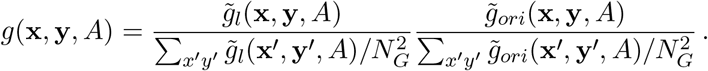

We then choose the number of connections 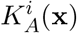 for the *i*^th^ cell at **x** from a Gaussian distribution with mean *K_A_*(**x**) = *pn_A_*∑*_y_ g*(**x**, **y**, *A*) and standard deviation *CV_K_* × *K_A_*(**x**), where *p* is a constant that is chosen such that the mean number of excitatory connections is similar to that of the unstructured model (for excitatory connections, ≈ 525 in our mouse models, ≈ 508 in our monkey models). We choose a total of 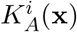 presynaptic neurons (excluding; autapses) by a weighted random sample without replacement, with weights *g*(**x, y**, *A*)/ (∑*_y′_ n_A_g*(**x, y**, *A*)) (choice function from python package numpy.random; if there is a potential autapse, it is removed from the sum in the denominator). The visual input to a cell of type A at **x** from a stimulus centered at **y** with orientation *θ_stim_* is 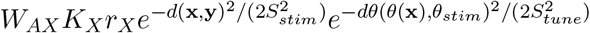. Here, *K_X_* = 0.1*N_X_* is the number of visual inputs to a cell and *r_X_* their firing rate. We set the input to have zero variance for simplicity. When we refer to visual-stimulus-matched cells, we mean cells with orientation tuning within 22.5° and receptive field within 0.6*dl* of the visual stimulus.

#### Network Simulations

Simulations of the network dynamics were performed in python. For each parameter set, we generated one realization of the network structure (adjacency matrices 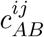, firing of the external population 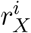 (generated from a Gaussian distribution with mean *r_X_* and standard deviation *σ_X_* = 0.2 *r_X_*), opsin expression 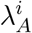 (generated from a lognormal distribution with mean *λ* and standard deviation *σ*_λ_) and simulated the network dynamics given by Eqs. (5) and (7) using the explicit Runge-Kutta method of order 5 (implemented with the function solve_ivp of the python package scipy.integrate). The network dynamics was run for a total simulated time of 200*τ_E_* = 4s for the unstructured model and 300*τ_E_* = 6s for the structured model and computed rates were stored every τ**i**/3 = 3.33ms. The firing rate of a cell was obtained by measuring its average rate over the simulation time (excluding the initial 10*τ_E_* = 0.2s for the unstructured model and 100*τ_E_* = 2s for the structured one.) To measure optogenetic responses, we simulated the same realization of network structure twice, once with 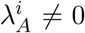 and once with 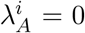. Parameters for all simulations are given in Tables 1 (unstructured models) and 2 (structured models).

**Table 1:** List of network parameters for unstructured models for Figures 4-6. Here and in Table 2 “FF input” means feedforward input, *i.e*. external (visual) input.

| Parameter | Symbol | Units | Figure 4<br>(weak, strong) | Figure 5 | Figure 6<br>(comb., mice, monk.) |
| --- | --- | --- | --- | --- | --- |
| Mean E connections per neuron | $K$ | | 2000 | 500 | 500 |
| Coeff. of Variation of In-Degree | $CV_K$ | | $(1-p)/\sqrt{K}$ | $(1-p)/\sqrt{K}$ | (0.015, 0.020, 0.007) |
| Connection probability | $p$ | | 0.1 | 0.1 | 0.1 |
| Number of excitatory neurons | $N_E$ | | $K/p$ | $K/p$ | $K/p$ |
| Ratio of I to E Population Size | $\gamma$ | | 0.25 | 0.25 | 0.25 |
| E time constant | $\tau_E$ | ms | 20 | 20 | 20 |
| I time constant | $\tau_I$ | ms | 10 | 10 | 10 |
| Synaptic efficacy | $J$ | mV | (0.01, 0.3) | (C) 0.6 | (0.14, 0.13, 0.44) |
| Ratio of I-to-E over E-to-E | $g_E$ | | 8 | (A,B,D) 8 | (4.2, 9.97, 4.58) |
| Ratio of I-to-I over E-to-I | $g_I$ | | 3 | (A,B,D) 3 | (3.24, 8.4, 2.73) |
| Ratio of E-to-E over E-to-I | $\beta$ | | 1 | 1 | (0.17, 0.33, 0.81) |
| Gain of the E population | $G_E$ | | from Eq. 6 | 1 | 1 |
| Gain of the I population | $G_I$ | | from Eq. 6 | 2 | 2 |
| Strength of FF input to E | $W_{EX}$ | | 3 | from Eq. 6 | from Eq. 6 |
| Strength of FF input to I | $W_{IX}$ | | (0.5, 2.5) | from Eq. 6 | from Eq. 6 |
| Mean FF input rates | $r_X$ | spk/s | 20 | (C) 10, (D) 17.97 | ([13.5 - 23.4] & [30.9 - 77.6], [7.24 - 13.8], [14.45 - 47.86]) |
| Std FF input rates | $\sigma_X$ | spk/s | $0.2r_X$ | $0.2r_X$ | 0 |
| Mean optogenetic input strength | $\lambda$ | mV | 20 | 20 | (1.9 & 5.5, 4.2, 2.5) |
| Std / mean of opsin expression | $\sigma_\lambda/\lambda$ | | 1 | (A,B) 2.15, (C) 1 | (1.69, 0.79, 6.6) |

**Table 2:**
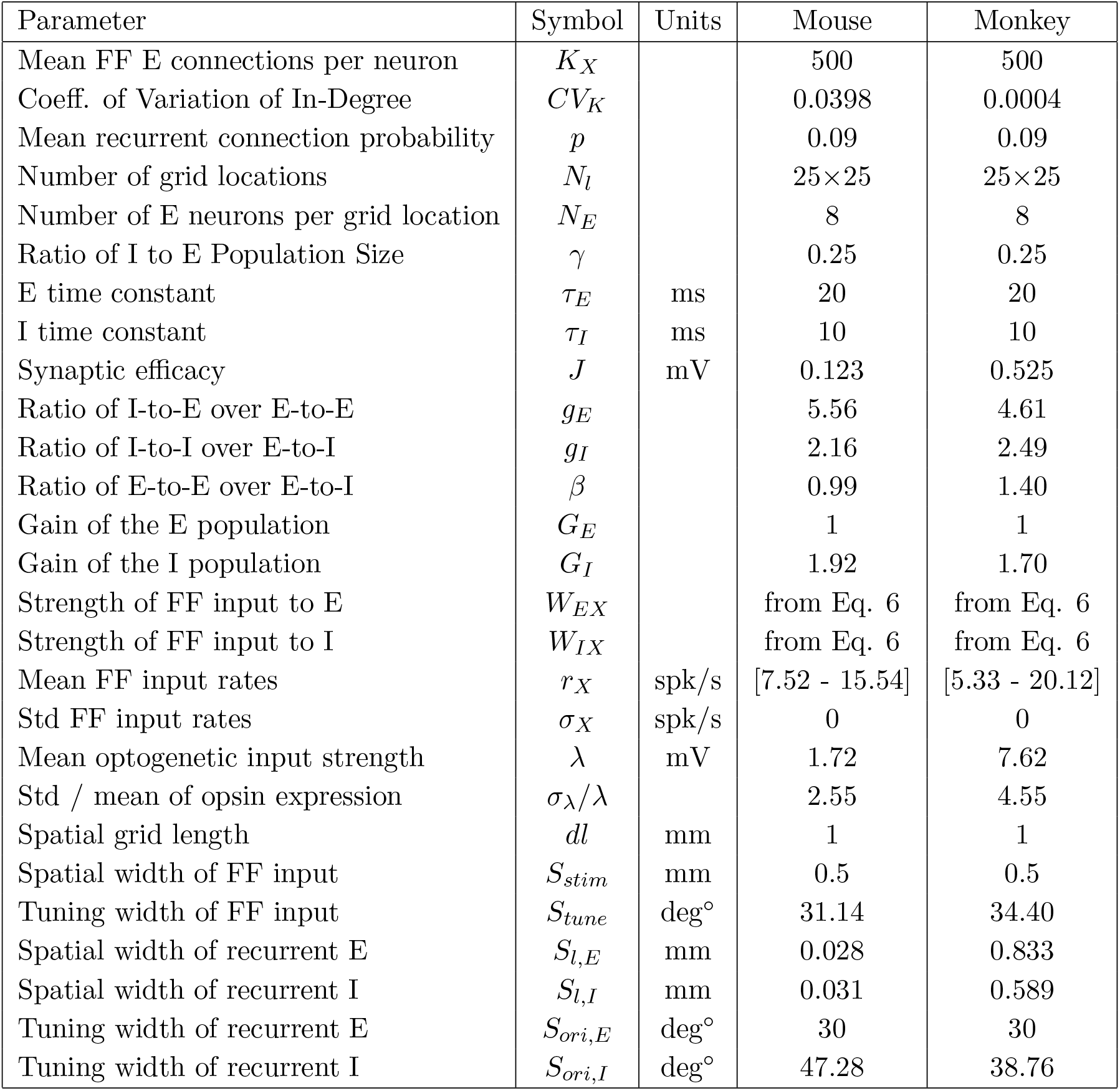
List of network parameters of best fit structured models for Figures 6 and S3.

### Fitting Procedure

#### Unstructured Model

Predictions of the model depend on a list of unknown parameters (indicated in what follows as Θ) related to: connectivity (*J, K, β, g_E_, g_I_, CV_K_*); external inputs (*G_E_, G_I_* and one value of *r_X_* per each contrast); and optogenetic inputs (*λ, σ_λ_*). To fit these parameters to experimental data, we assumed *G_I_* = 2*G_E_*, consistent with the fact that in vivo, I firing rates are typically about twice E firing rates (e.g. (Sanzeni et al., 2020)). This choice is motivated by the fact that experimental recordings do not distinguish cell types; this limitation makes our model under-constrained by the data (since the same average firing could be obtained by an infinite combination of firing rates of E and I cells). Without loss of generality, we assumed *G_E_* = 1 (any other choice can be obtained by rescaling *r_X_*) and *K* = 500 (any other choice can be obtained by rescaling 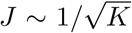). Moreover, we took *σ_X_* = 0, and controlled variability in the external input by varying *CV_K_*.

With the above assumptions, model parameters were inferred as follows. First, we measured optogenetic responses in a large number (770195) of network simulations; each one obtained assuming model parameters generated randomly and uniformly in the intervals

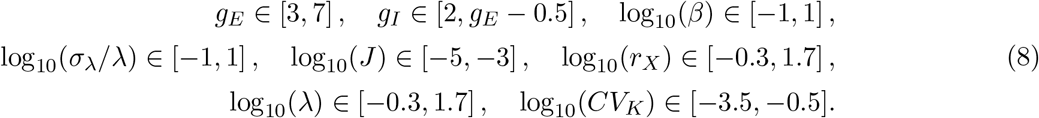

Starting from the results of these simulations, we trained a multilayer perceptron (MLPRegressor from python package sklearn.neural_network (Pedregosa et al., 2011); specifics were activation=ReLU, hidden_layer_sizes=(100, 150, 50),)) to predict the mapping *f* (Θ) between model parameters (Θ) and moments of the optogenetic response (*r*, Δ*r, σ_r_, σ*_Δ*r*_, Cov(*r*, Δ*r*)). The learned multilayer perceptron was able to predict the outcome of simulations with high accuracy (with 20% of held-out simulations, the crossvalidated performance had *R*^2^ = 0.924). We then fitted model parameters Θ to data by minimizing the squared error between predictions given by *f* (Θ) and moments of the optogenetic responses measured in experiments (least_squares function from python package scipy.optimize). In the fits, we excluded simulation parameters leading to strongly chaotic dynamics by imposing an additional penalty in the least squared error for networks whose predicted autocorrelation time was smaller than 1 second. Specifically, all moments of the network response for which the predicted autocorrelation time was smaller than 1 second were set to 10^10^; this prevented the least square optimization algorithm from exploring regions of the parameter space that lead to strong chaotic dynamics.

#### Structured Model

In order to fit the structured model, we allowed *G_I_* to vary since the spatial structure often causes the network to be more inhibition dominated than the unstructured model. As for the unstructured model, we measured optogenetic responses in a large number (484848 with salt-and-pepper maps and 508719 with smooth maps) of network simulations; we generated parameters randomly and uniformly in the following intervals

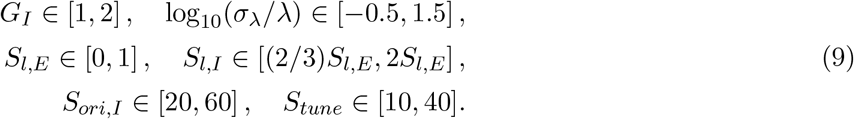

In addition, we excluded parameters where the parameterization from Eq. (6) would give negative mean external input to any population. We fixed *S_ori,E_* = 30° and *S_stim_* = 0.5*dl*. All other parameters not listed above are generated with the same intervals as for the unstructured model. For each simulation, we also generate a different random orientation map.

Instead of training a multilayer perceptron to predict the mapping between model parameters and moments as in the unstructured model, we looked for simulations that already had a small squared error between the simulated predictions and the moments from the experimental data at any contrast. For these models we then resimulated them, averaging moments across four different orientation maps, and lastly if the moments still robustly fit the data after averaging then we calculated the moments at various *r_X_* in order to find a mapping between *r_X_* and contrast that minimized the squared error (least_squares function from python package scipy.optimize).

#### Mean Field Theory for the Unstructured Model

For *N_B_* ≫ 1 in the unstructured model, 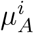 and 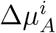 in Eqs. (5) and (7) are well approximated by correlated random Gaussian variables with

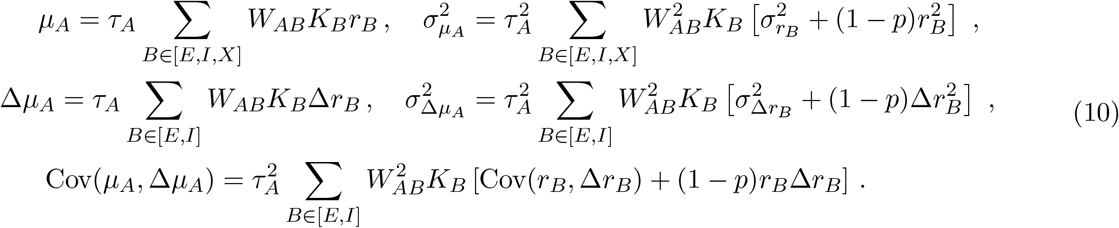

In Eq. (10), *μ_A_* (Δ*μ_A_*) and *σ_μ_A__* (*σ*_Δ*μ_A_*_) are the mean and variance of 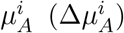. The covariance Cov(*μ_A_*, Δ*μ_A_*) is computed starting from

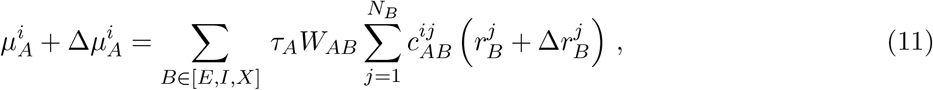

applying the identity Var(*X + Y*) = Var(*X*)+Var(*Y*) + 2Cov(*X, Y*) to both sides and using the central limit theorem. Note that optogenetic input affects firing differently than external and recurrent inputs. In fact, firing depends on the whole distribution of 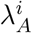 but only on the first two moments of external and recurrent inputs.

If the network dynamics settle in a stable fixed point, the statistics of currents and rates in the network are related by

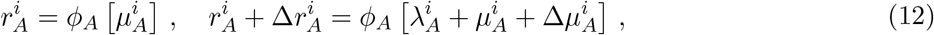

and moments of the rates can be computed by self-consistently solving a set of ten equations

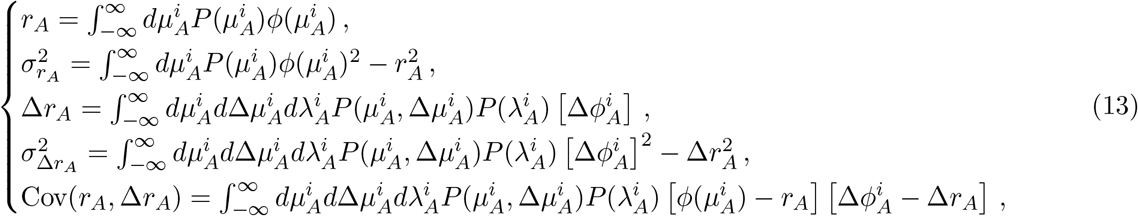

with 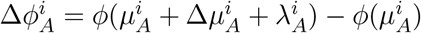.

If the fixed point of the network dynamics is unstable, rates and currents become time-dependent. One can again use a Gaussian ansatz, in which the total inputs at different times 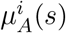 and 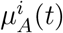 are jointly Gaussian with means *μ_A_*(*s*), *μ_A_*(*t*) and covariance 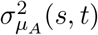. Analogously, rates at different times 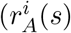 and 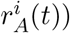 are jointly Gaussian with means *r_A_*(*s*), *r_A_*(*t*) and covariance 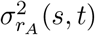. At any given pair of time points *t* and *s*, the moments of rates and currents are related by Eq. (10). The temporal evolution of the moments can be computed using standard techniques (Sompolinsky et al., 1988). In particular, indicating with 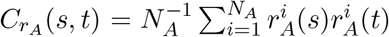 the autocorrelation function of the rates of population *A* at times *s* and *t*, we can define the autocovariance function of the rates as

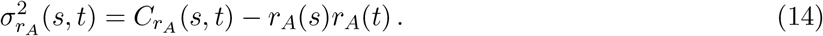

By multiplying Eq. (7) at times *s* and *t*, averaging over cells in population *A*, and rearranging terms we see that *C_r_A__*(*s,t*) evolves in time according to

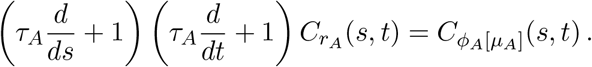

We can repeat the above steps to derive equations for the evolution in time of all five order parameters per population of neurons (mean, autocovariance, and cross-covariance of rates without and with optogenetic stimuli). Defining

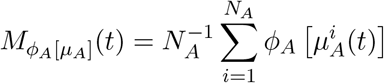

and

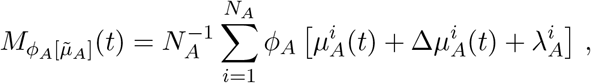

the order parameters evolve in time according to

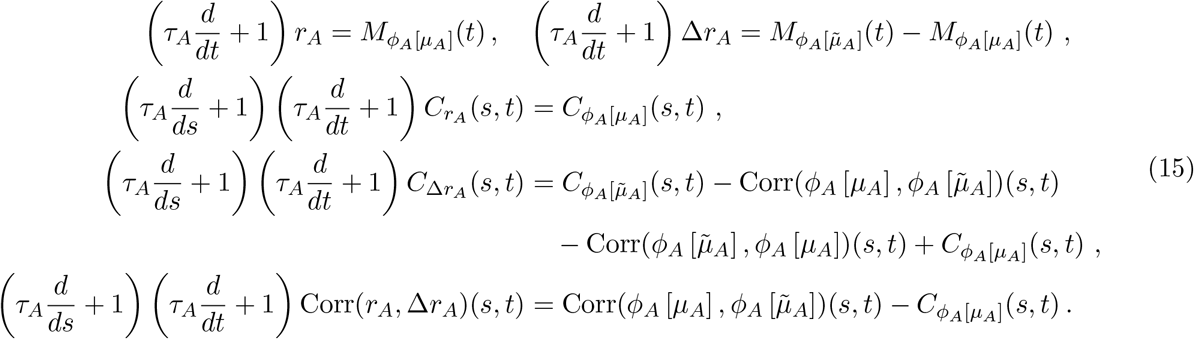

#### Numerical Solution of Mean Field Equations

We solve for the stationary statistics of the chaotic network by explicitly integrating Eq. (15) in time until the order parameters converge to a steady state. We start with initial values for *r_A_*(0) and Δ*r_A_*(0) as well as initial functions for *C_r_A__*(*s*, 0) = *C_r_A__*(0, *t*), *C*_Δ*r_A_*_(*s*, 0) = *C_r_A__*(0, *t*), and Corr(*r_A_*, Δ*r_A_*)(*s*, 0) = Corr(*r_A_*, Δ*r_A_*)(0, *t*). The means *r_A_*(*t*) and Δ*r_A_*(*t*) are integrated with the usual explicit RK4 method, but the autocovariance and cross-covariance functions are integrated on a grid with an explicit Euler-like method derived using forward finite differences. For example, when integrating *C_r_A* (s, t) we can express *C_r_A__* (*s + dt, t + dt*) in terms of *r_A_*(*s*), *r_A_*(*t*), *C_r_A__* (*s, s*), *C_r_A__*(*t, t*), *C_r_A__*(*s, t*), *C_r_A__*(*s+dt, t*), and *C_r_A__* (*s,t+dt*). Assuming that we have solved for the order parameters at all time points (*s* ≤ *t*_0_, *t* ≤ *t*_0_), we can explicitly solve for the order parameters at time points (*s* ≤ *t*_0_ + *dt, t*_0_ + *dt*) and (*t*_0_ + *dt, t* ≤ *t*_0_ + *dt*). Furthermore, we can ignore portions of the grid for |*s* – 1| > *T* for some *T* that is much longer than the width of the steady state autocorrelation function in order to reduce the computation time required for the integration.

A useful simplification of the equations governing the evolution of the order parameters is to calculate quantities such as *C*_*ϕ_A_*[*μ_A_*]_(*s, t*) by setting the mean and variance of 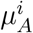 at times *s* and *t* equal to the mean and variance at time max(*s, t*). While this change increases the required integration time for the order parameters to reach the steady state, it reduces the number of dependent variables needed to calculate quantities like *C*_*ϕ_A_*[*μ_A_*]_(*s, t*) from five to three. With only three dependent variables, it becomes feasible to pre-calculate these autocorrelation functions on a 3D grid and interpolate between the pre-computed points in order to accelerate the integration. In addition, we can precompute quantities such as *M*_*ϕ_A_*[*μ_A_*]_(*t*), which only depends on the mean and variance of the inputs, on a 2D grid and interpolate them as well.

The cross-correlation function 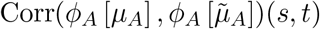 cannot be pre-computed and interpolated since the means and variances of the cross-correlated variables are not equal. However, since we have already interpolated *M*_*ϕ_A_*[*μ_A_*]_(*t*) and 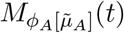 we can efficiently calculate the cross-correlation function from a single Gaussian integral.

Indicating with 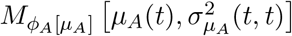 and 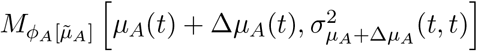 the precomputed mean rates as a function of the means and variances of the inputs, we can express the crosscorrelation function as

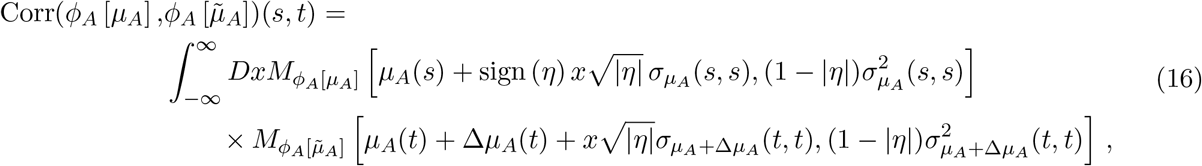

with the Pearson correlation coefficient *η* defined as

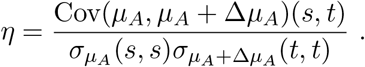

## Acknowledgments

This work was supported by the NIMH Intramural Research Program and by NIH BRAIN U01 NS108683 (to N.B., M.H.H., K.D.M. and J.H.R.). A.S., A.P., T.H.N. and K.D.M. were partially supported by the Gatsby Charitable Foundation GAT3708 and by NSF Grant DBI-1707398. A.P. and K.D.M. were partially supported by NIH BRAIN U19 NS107613. A.P. would like to acknowledge the support of the Swartz Foundation Fellowship for Theory in Neuroscience 2019-4 and 2020-6. T.H.N. would like to acknowledge the support by NIH grant 5R01EY029999. J.J.N was funded, in part, by NIH grant R01-EY021827. J. Luo was supported, in part, by the Fiona and Sanjay Jha Chair in Neuroscience. This work used the computational resources of the Duke Compute Cluster.

## Supplementary Information

### Theoretical results are preserved with different transfer functions

**Figure S1:**
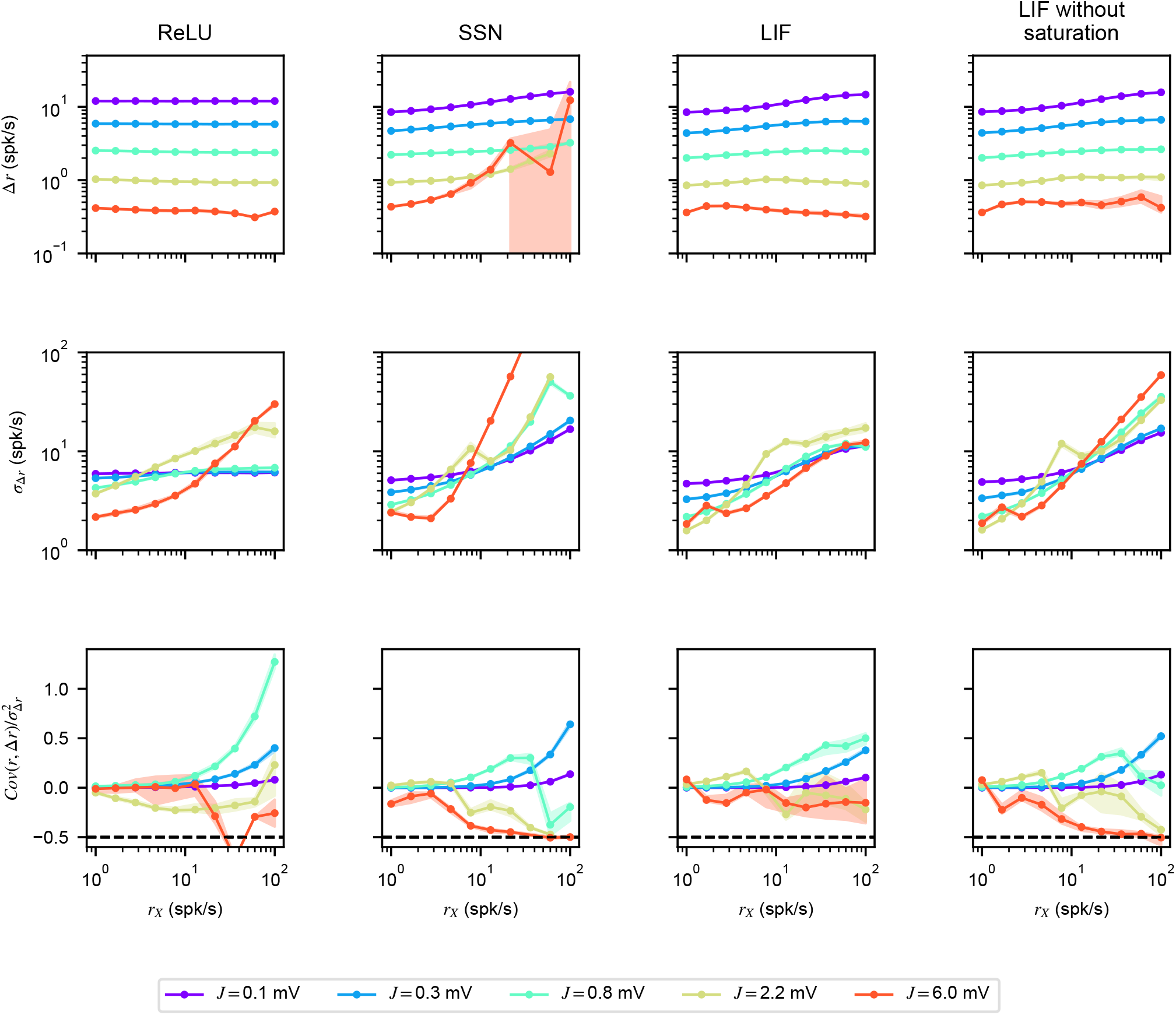
Neuronal reshuffling emerges in network models with different single transfer functions. Optogenetic responses obtained in numerical simulations of inhibitory networks (dynamics given determined by Eqs. (5) and (7), without E cells and with optogenetic stimuli targeting I cells) as a function of *J* (colors) and *r_X_*, for different choices of the single neuron transfer function *ϕ_A_* (titles). Plots are obtained averaging over 20 realizations of the network models (dots represent means, shaded regions are mean±sem). Transfer function used were: *ϕ*(*x*) = *a* [*x*]_+_ (ReLU, *a* = 1.5 mV^-1^); 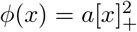 (SSN, *a* = 0.05 mV^-2^); *ϕ*(*x*) as in Eq. (3) with *τ_rp_* =2ms (LIF) or *τ_rp_* = 0 (LIF without saturation).

### Chaos is not required for negative covariance to appear

**Figure S2:**
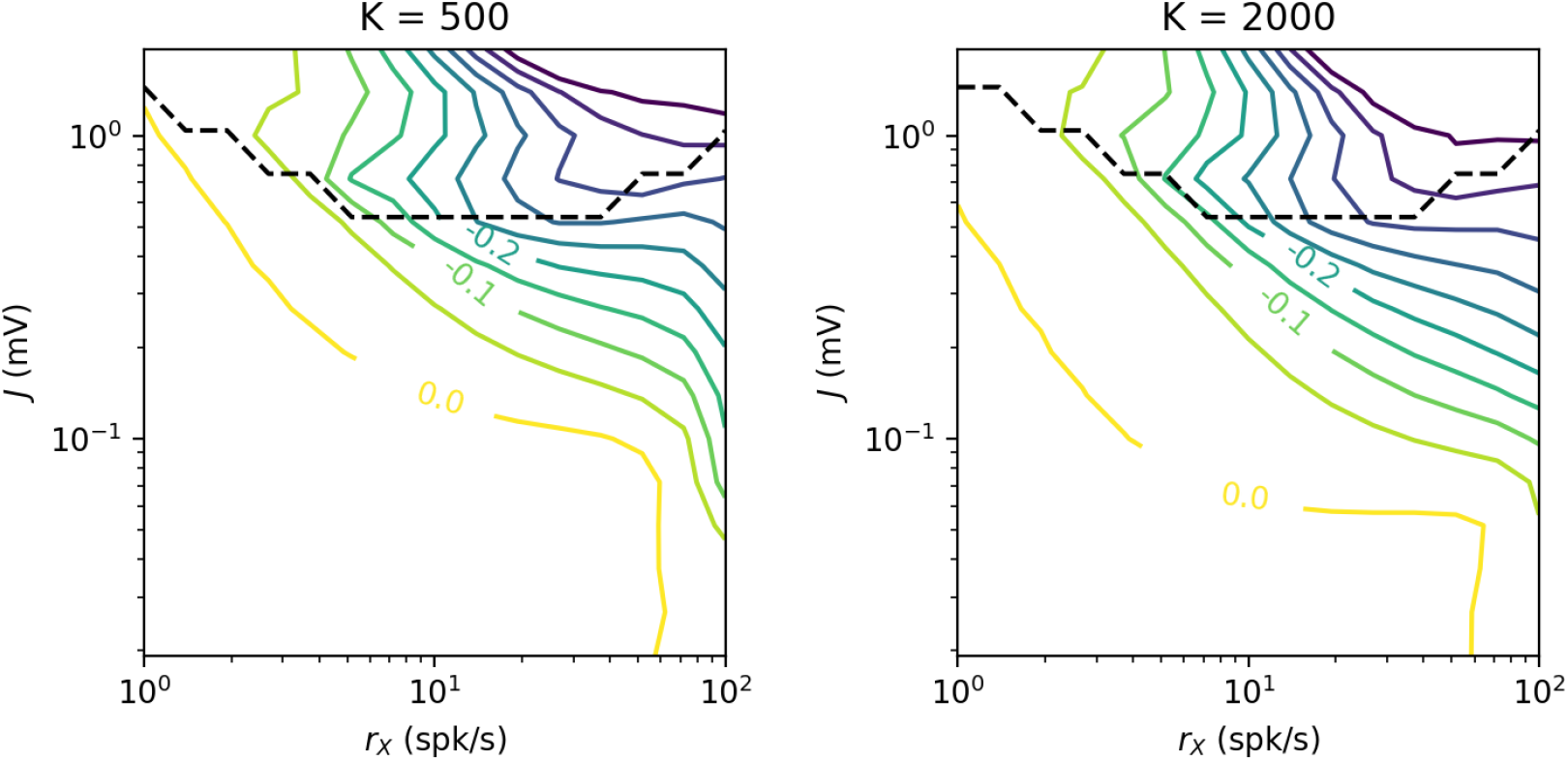
Phase diagram of chaos and negative covariance. Colored lines indicate contours of constant normalized covariance between 0 and −0.5 as a function of the external input *r_X_* and recurrent synaptic efficacies *J*. The dashed line indicates the transition to chaos; Below the dashed line the baseline system is stationary, and above the line the network is chaotic. In order to investigate the effect of finite network size, the left plot is calculated with *K* = 500 while the right plot is calculated with *K* = 2000. Increasing *K* tends to increase the magnitude of the normalized covariance while decreasing the region of chaos. All values are calculated from mean field theory. Network parameters are identical to those used in Figure 5 panels **A-B**.

### Structured Connectivity Reshuffles Visual-Stimulus-Matched Rates Even with Large Optogenetic Input

**Figure S3:**
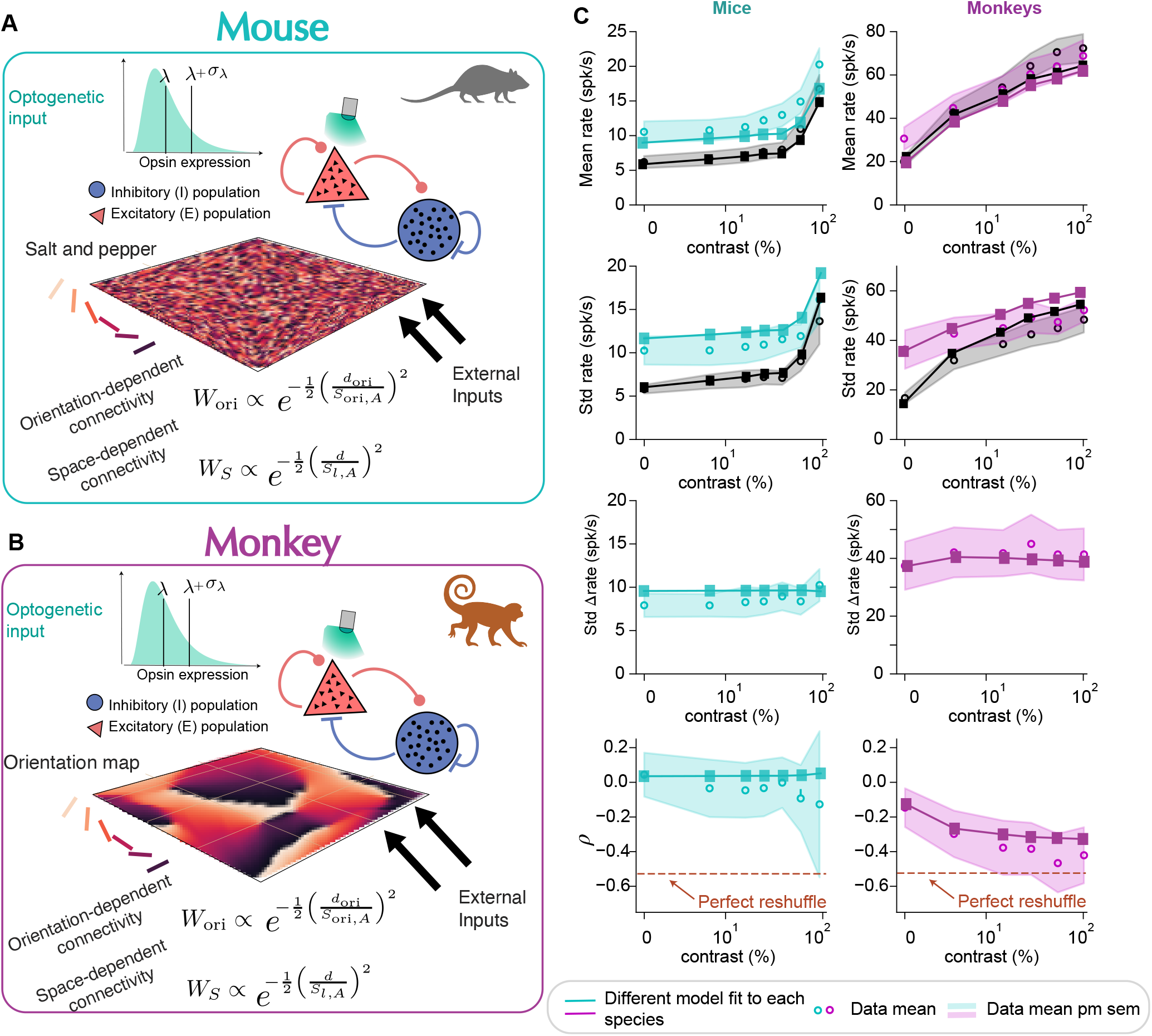
Structured models capture the data. **A** Structure of the mouse model. The model consists of a grid of 25×25 spatial locations. Each spatial location contains a total of 10 neurons, 8 excitatory and 2 inhibitory, to match the size of the unstructured networks. The orientation preference of each cell has a salt-and-pepper arrangement in the 2D space, the external inputs are a Gaussian function of the difference between their preferred orientation and the orientation of the stimulus. The connectivity between two cells is proportional to the product of two Gaussian functions, one of the difference between the preferred orientation of the cells, *W_ori_*, and another of their spatial location, *W_S_*. As in the structure-less model described Fig. 4, external inputs target both inhibitory and excitatory cells while optogenetic inputs target only excitatory cells. **B** Structure of the monkey model. Idem (A) but the spatial arrangement of the orientation preference is given by an aperiodic map. Parameters of these fits can be found on Table 2. **C** Comparison between data (circles represent means, shaded regions are mean±sem) and network models (full lines with squares) as described in A for the mouse and in B for the monkey. In the mouse, all cells were taken into account in order to perform the averages, as was done in the data. For the monkey, only visual-stimulus-matched cells were used to fit the data.

**Figure S4:**
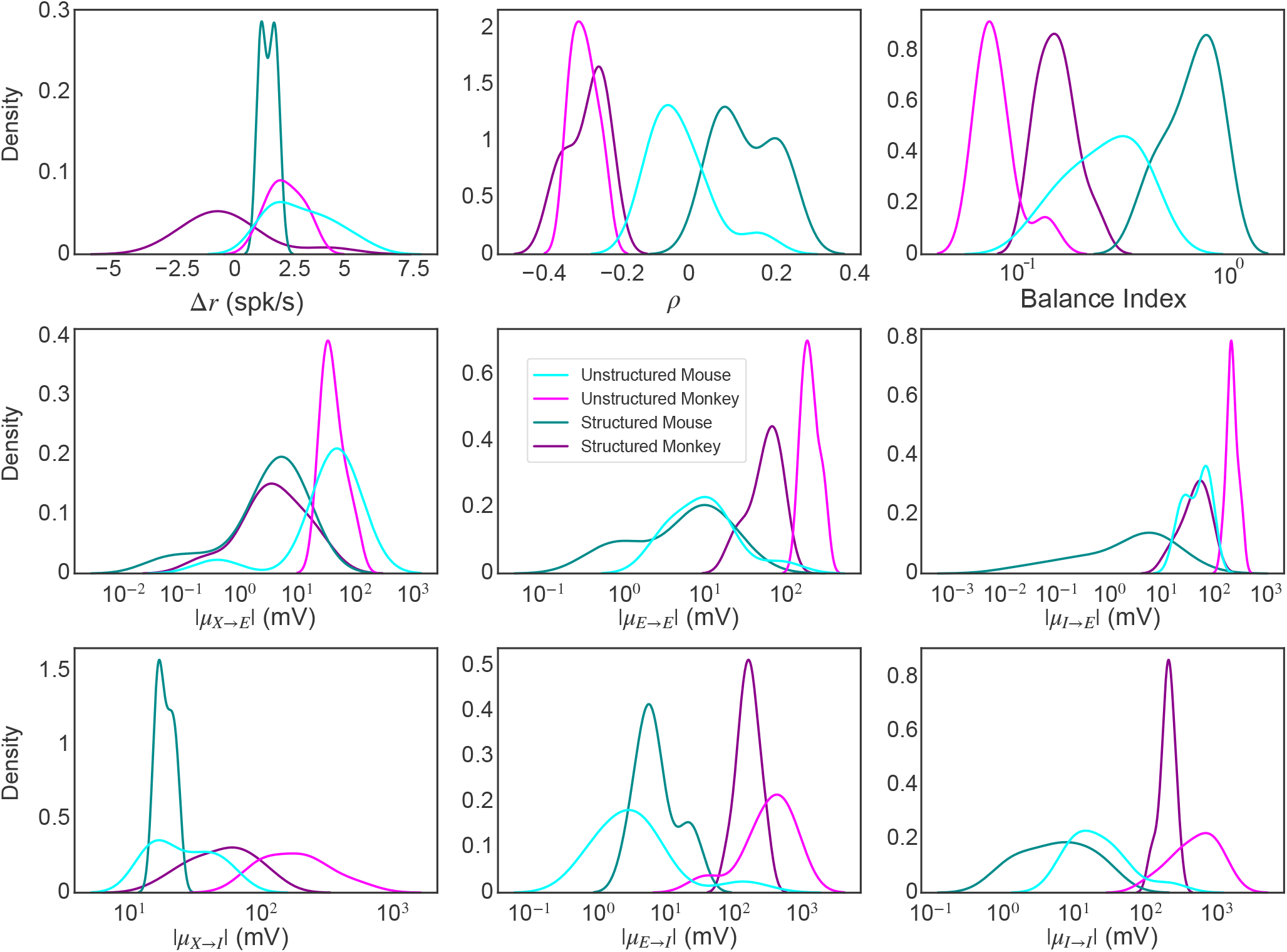
Unstructured models require tighter balance to fit the data. Kernel density estimation plots of Δ*r*, normalized covariance *ρ*, balance index, and absolute mean inputs *μ_A→B_* from inputs of type A to cells of type B, for unstructured and structured models that best fit experimental data for each animal at the highest contrast. All quantities are calculated from the entire network except for structured monkey models, where we restrict our analysis to only visual-stimulus-matched cells as we did for the fitting procedure. Only structured models with quasi-periodic maps (fit to experimental results for the monkey) are found to have negative Δ*r*, suggesting an alternative mechanism for the small change in mean response compared to unstructured networks. Structured networks tend to have looser balance (higher balance index) and smaller absolute mean inputs compared to unstructured networks fit to the same animal data (except for the E → I input for the mouse). Models were selected by taking all previously simulated optogenetic responses and repeating the fitting procedure used for the structured models in order to find model parameters that had the least squared error between simulated responses and experimental data at all contrasts. We then chose cutoffs for the squared error to select about 10 of the best models for each case, and plotted kernel density estimations using results at the highest contrast.

**Figure S5:**
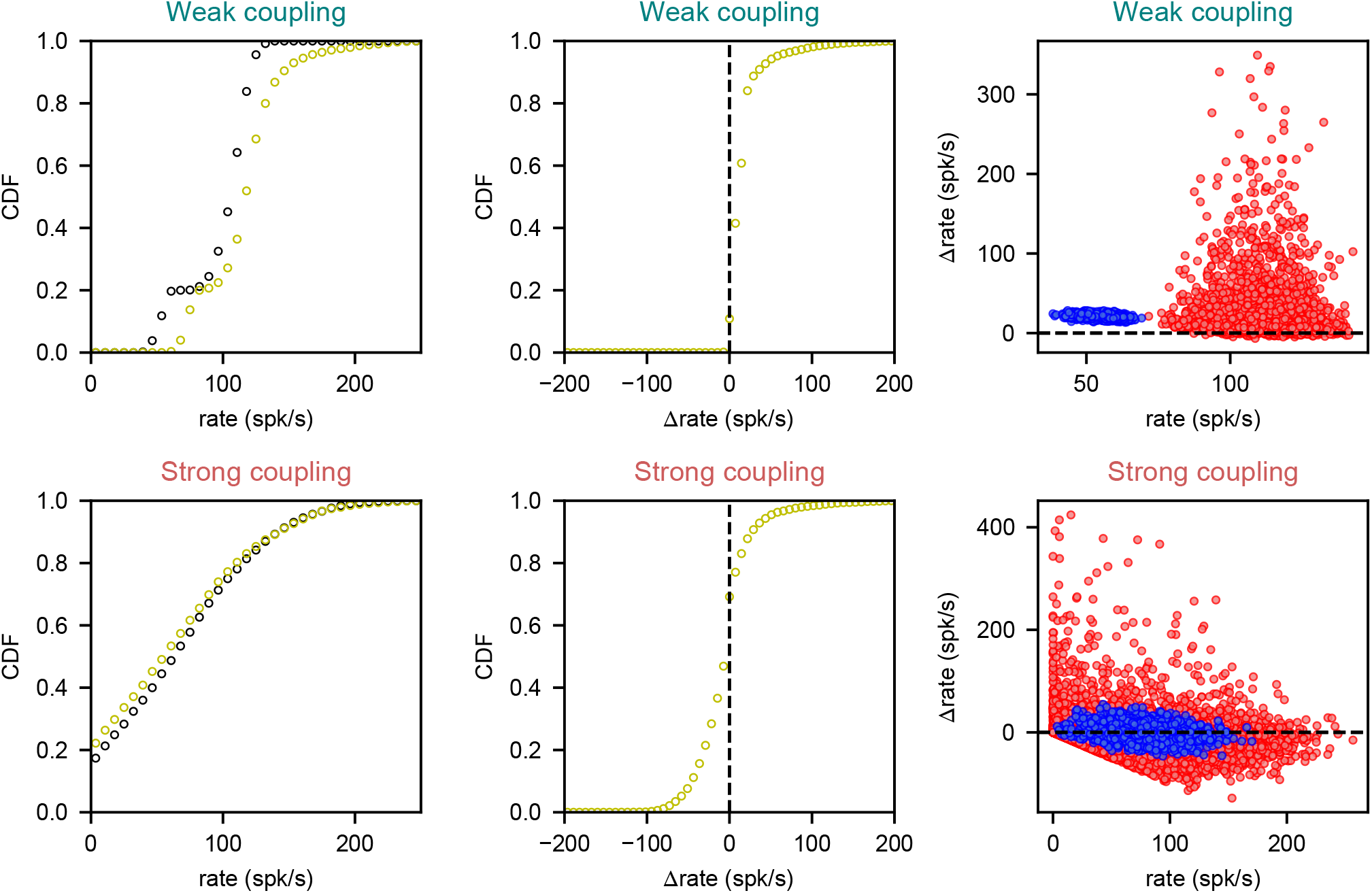
Strong coupling is also necessary for reshuffling in structured networks. Panels are arranged as in Figure 4 **B-G**, but instead show the firing rates of visual-stimulus-matched cells in a structured network with an quasi-periodic map. With weak coupling, optogenetic stimulation tends to excite most cells whereas with strong coupling, the distribution of rates is not significantly altered by optogenetic input. The normalized covariances *ρ* (for E, I, and E+I cells, respectively) are −0.03, −0.64, and −0.03 for the weakly coupled network and −0.28, −0.44, and −0.29 for the strongly coupled network. The Pearson correlation coefficients are −0.08, −0.32, and −0.04 for the weakly coupled network and −0.22, −0.26, and −0.22 for the strongly coupled network. All parameters except the synaptic efficacy *J* are taken from the best structured fit to the monkey data at the highest contrast (see Table 2). In the top row, the best fit synaptic efficacy is decreased by a factor of six, whereas in the bottom row the synaptic efficacy is taken from the best fit.

